# How Much Can AI Virtual Cells Gain from Unified Perturbational Modeling?

**DOI:** 10.64898/2026.10.25.761625

**Authors:** Zongting Wei, Yuedian Ye, Shuguang Cui, Wenjing Ma, Yuning You

## Abstract

Predicting how individual cells respond to perturbations is a central task in artificial intelligence virtual cell (AIVC) modeling. However, integrating perturbation datasets is challenging because they differ in measured gene sets, perturbation types and cellular contexts. Existing AIVC predictors are typically developed for specific settings, limiting their applicability across heterogeneous datasets. Here, we investigate the extent to which AIVC predictors can benefit from unified perturbational modeling (UPM), a novel paradigm designed to integrate heterogeneous Perturb-seq datasets into a single predictive model. To this end, we introduce three unification modules into a variational autoencoder (VAE)-based AIVC predictor to address these sources of heterogeneity: gene-channel alignment, shared perturbation encoding, and joint embedding of gene expression and identity. We systematically evaluate and validate the individual contributions of these modules. Building on UPM, we assemble a dataset of 110 million cells spanning over 22,000 genes, four perturbation types and 76 cellular contexts, and use 89.7 million cells from 71 contexts to train a unified AIVC predictor with more than one billion parameters. Across genetic perturbation benchmarks, UPM achieves the highest PES and gene-wise response correlation among the evaluated models. It also improves all four evaluation metrics over State on the five held-out Tahoe-100M cell lines and over Stack on the cytokine benchmark. UPM also improves combinatorial perturbation prediction relative to specialized predictors. In the HSC application, our model achieves a 40.0% improvement in accuracy on a downstream task involving the recapitulation of effects induced by the knockdown of hepatic stellate cell (HSC)-related targets. Our implementation is openly available at https://github.com/youlab2025/UPM.

## Introduction

Predicting how individual cells respond to perturbations is a central task in artificial intelligence virtual cell (AIVC) modeling [1, 2]. By predicting cellular responses to perturbations *in silico*, AIVC models have the potential to prospectively guide experimental design and decision-making in wet-laboratory studies [3]. The rapid accumulation of Perturb-seq datasets, which measure cellular responses to perturbations at single-cell resolution, has enabled the development of increasingly capable perturbation effect prediction models [4, 5, 6].

However, Perturb-seq datasets differ substantially in measured gene sets, perturbation types and cellular contexts [7]. Existing AIVC models remain specialized for particular subsets of these data [8]. Their modeling strategies and scope are reviewed in Appendix A. Integrating these heterogeneous resources could broaden the applicability of AIVC models to therapeutic target prioritization and drug screening across diverse cellular contexts.

In this paper, we investigate whether a unified perturbation effect predictor can be developed by integrating heterogeneous Perturb-seq datasets, and to what extent AIVC models can benefit from such unification. A unified approach could offer three potential benefits. First, in terms of applicability, a unified model could accommodate a substantially broader range of prediction scenarios, thereby expanding the practical utility of AIVC models. Second, in terms of predictive performance, joint training on heterogeneous Perturb-seq datasets could leverage complementary information and synergies across data sources, potentially improving prediction accuracy within individual perturbation types and cellular contexts [9, 10, 11]. Third, in terms of representation learning, integrating diverse perturbation types and cellular contexts could reshape the learned latent representations by uncovering previously unexplored relationships among perturbations and cellular states, potentially yielding more generalizable representations [12].

To this end, we introduce three unification modules into a variational autoencoder (VAE)-based perturbation effect predictor to address heterogeneity in gene measurements, perturbation types, and cellular contexts across Perturb-seq datasets. We term this framework unified perturbational modeling (UPM). (i) Gene-channel alignment: We align gene channels across datasets using their Ensembl identifiers and construct a unified gene set comprising the union of all measured genes, enabling the model to accommodate heterogeneous gene coverage and make predictions across different gene sets. (ii) Shared perturbation encoding: We learn perturbation representations specific to each perturbation type and map them into a shared latent space through a common projection module. (iii) Joint embedding of gene expression and identity: We jointly embed gene expression values and their corresponding gene identities into a cell-state latent representation, enabling the model to accommodate different perturbation types, represent and process cells from different cellular contexts, and predict their effects within a VAE-based model.

We systematically assess the contributions of these three modules by comparing the unified model with its specialized counterparts. By training nine model configurations and benchmarking their performance across four metrics, we demonstrate two key findings. (i) Gene channel unification preserves predictive performance relative to models specialized for particular gene subsets, despite the increased processing and prediction complexity introduced by the union of measured gene sets. (ii) Unification across perturbation types and cellular contexts consistently improves mean-response prediction, with additional gains in perturbation discrimination measured by the perturbation effect score (PES) across cellular contexts and variable effects across perturbation types.

Building on UPM, we train a unified AIVC model with over one billion parameters on 89.7 million cells from 71 cellular contexts drawn from the full dataset. Across genetic perturbation benchmarks, UPM achieves the highest PES and gene-wise response correlation among the evaluated models [13, 14]. It also improves all four evaluation metrics over State on the five held-out Tahoe-100M cell lines and over Stack on the cytokine benchmark [15, 16]. UPM also improves combinatorial perturbation prediction relative to specialized predictors. Furthermore, we apply our model to a hepatic stellate cell (HSC) dataset [17, 18], achieving a 40.0% improvement in accuracy on a downstream task assessing the recapitulation of effects induced by the knockdown of HSC-related targets.

## Results

### Unified perturbational modeling enables one AIVC model to integrate Perturb-seq data from 110 million cells spanning over 22,000 genes, four perturbation types, and 76 cellular contexts

#### UPM integrates heterogeneous perturbation datasets through three unification modules within a VAE-based architecture

Given an unperturbed cell state and one or more perturbation inputs, UPM predicts perturbed single-cell expression in a shared coordinate space of 22,768 genes (Figure 1A). The three modules provide gene-channel alignment, shared perturbation encoding, and joint embedding of gene expression and identity. The VAE-based architecture combines a cell-state encoder, a shared perturbation encoder, and a cell-state decoder (Figure 1B; Appendix B.1). A wide multilayer perceptron encodes the unperturbed expression profile. Projections specific to each perturbation type map perturbation features to a common dimensionality before processing by the shared perturbation encoder. For combined perturbations, the constituent embeddings are averaged. The cell-state and perturbation embeddings are concatenated and passed through separate networks to parameterize the mean and log-variance of a diagonal Gaussian latent distribution. During inference, the decoder transforms the latent mean into an expression change that is added to the unperturbed input. The four perturbation types comprise CRISPRi, CRISPRa, drug and cytokine (Figure 1A). Drug dose is incorporated into the drug perturbation representation. This architecture predicts responses to individual perturbations and combinations within and across perturbation types.

**Figure 1:**
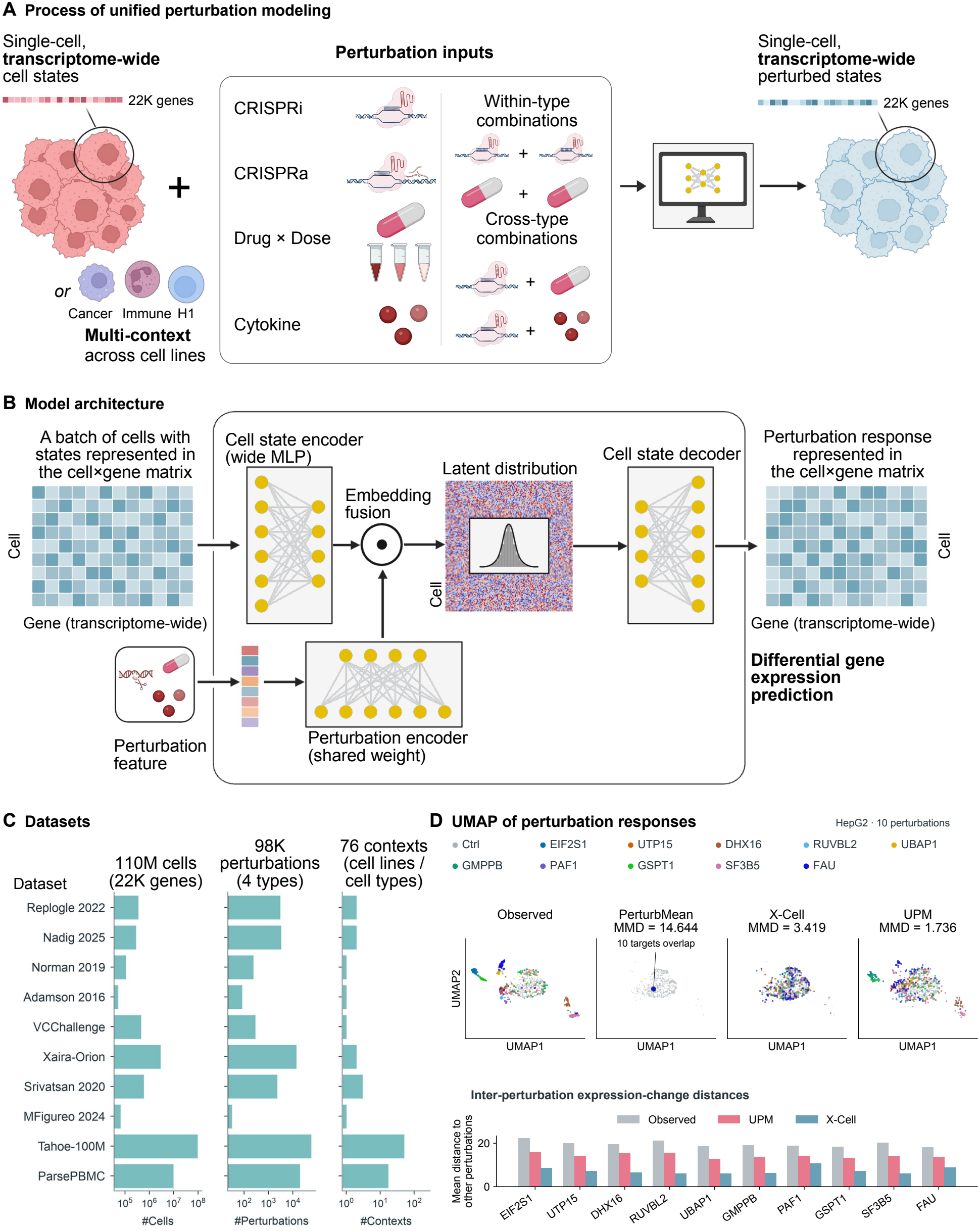
UPM unifies gene channels, perturbation types and cellular contexts across a resource of 110 million cells. **(A)** Prediction workflow. UPM combines unperturbed cell states with CRISPRi, CRISPRa, drug or cytokine inputs to predict responses to individual perturbations and combinations within and across perturbation types. Drug representations incorporate dose information. **(B)** VAE-based encoder-decoder architecture incorporating gene-channel alignment, shared perturbation encoding, and joint embedding of gene expression and identity. Cell-state and perturbation representations parameterize a latent distribution, and the decoder predicts expression changes that are added to the unperturbed input. **(C)** The complete resource spans over 22,000 genes, four perturbation types and 76 distinct cellular contexts (58 cell lines and 18 PBMC cell types). Bars show context counts within each dataset; the total is deduplicated across datasets. Following filtering and held-out splits, the training collection contains 89.7 million cells and 71 contexts (53 cell lines and 18 PBMC cell types), represented using a shared set of 22,768 genes. **(D)** Observed and predicted single-cell expression distributions for ten perturbations selected from the 258 held-out HepG2 test perturbations based on differences in observed responses across all 9,191 measured genes (Appendix B.4). Predictions are projected into a common PCA–UMAP embedding fitted to observed cells and controls; colors identify perturbations and controls. The Observed, UPM and X-Cell [14] panels each contain 542 perturbed cells and 512 controls, with cell counts matched across panels for each perturbation. UPM averages predictions from two unified model configurations with equal weights. PerturbMean assigns the same mean response to every perturbation, producing overlapping point predictions. Mean MMD, which measures differences between predicted and observed cell distributions, is 1.736 for UPM, 3.419 for X-Cell and 14.644 for PerturbMean. Bars below the UMAP plots show each perturbation’s mean Euclidean distance to the other nine perturbations, calculated from their mean expression-change vectors.

#### The collected perturbation resources comprise approximately 110 million cells across 76 distinct cellular contexts

After training exclusions and held-out splits, the final expression-training collection contains 89.7 million cells across 71 cellular contexts, with expression profiles aligned to 22,768 genes (Figure 1C). Cellular contexts are counted as distinct cell lines or PBMC cell types across datasets, with donors reported separately. The five contexts excluded from training are held-out Tahoe-100M cell lines. Genetic perturbation data come from Adamson, Replogle, Nadig, Norman, Virtual Cell Challenge 2025 and X-Atlas/Orion [5, 6, 19, 20, 21, 22]. Chemical and cytokine data come from sci-Plex 3, SciPlexCombo, Tahoe-100M and the Parse PBMC cytokine resource [23, 24, 25, 26]. Selected target genes, Norman and SciPlexCombo combinations, drug–dose conditions, and donors are held out for validation or testing. No gene combinations enter the training collection. Complete inventories, counting definitions and split details are provided in Appendix B.2.

#### Unified modeling substantially enhances AIVC models’ ability to capture single-cell perturbation-response distributions

Single-cell distribution prediction was assessed using maximum mean discrepancy (MMD), and perturbation discrimination using perturbation effect score (PES). ΔPearson and ΔPert *r* quantify control-centered mean-response correlations across genes and perturbation conditions, respectively. Lower MMD and higher values of the other metrics indicate better performance (Appendix B.3). UPM predictions combined two unified model configurations with equal weights. Across all 9,191 measured genes in the ten displayed HepG2 perturbations, UPM reduced MMD by 49.2% relative to X-Cell [14], with higher PES and both mean-response correlations (Figure 1D). UPM predictions showed closer visual agreement with observed cells in the UMAP projections and more closely reproduced the observed mean distances between perturbation responses than X-Cell for all ten perturbations. Evaluation across all 258 held-out HepG2 perturbations also showed improvements over Gene-editing-only, with 8.6% lower MMD and higher PES and both mean-response correlations.

### Evaluation of UPM across training settings and perturbation benchmarks. Together, UPM achieves SOTA performance across genetic, drug, and cytokine benchmarks

#### Perturbation-effect prediction performance is evaluated across diverse unseen perturbations and cellular contexts using both distributional and bulk-level metrics

Models with different supervision over gene channels, perturbation types and cellular contexts were compared to assess the contribution of each dimension of unification (Figure 2A–C). Evaluation groups the four perturbation types into three categories: genetic perturbations comprising CRISPRi and CRISPRa, chemical perturbations, and cytokine stimulation. Each perturbation condition is specified by the intervention and cellular context, together with the dose for drug and cytokine treatments. Percentage changes are calculated relative to the specified comparator. The UPM models in Figure 2A–C correspond to their respective training-scope comparisons, whereas the external benchmarks in Figure 2D–F use the same final unified model.

**Figure 2:**
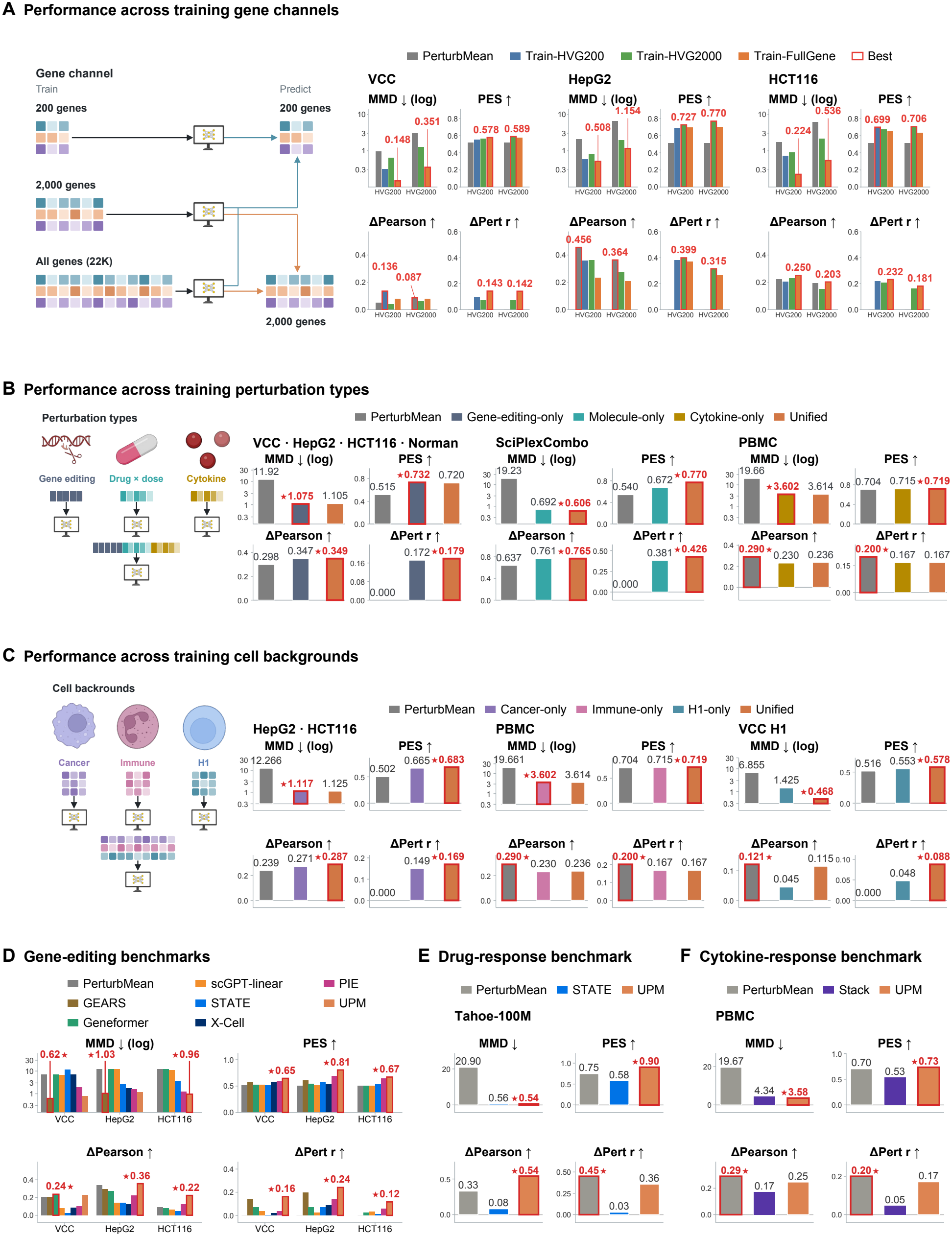
Evaluation of UPM across training settings and perturbation benchmarks. A–C compare models trained with different gene supervision, perturbation types or cellular contexts. D–F evaluate the same final unified UPM model across external benchmarks. **(A)** Comparison of HVG200, HVG2000 and full-gene supervision on matched evaluation gene panels in VCC, HepG2 and HCT116. All three training settings are evaluated on the HVG200 panel; HVG2000 and full-gene training are also compared on the HVG2000 panel. Retained gene counts are reported in Table S1, and training and checkpoint details in Appendix B.4. **(B)** Unified training compared with Gene-editing-only on genetic benchmarks, Molecule-only on SciPlexCombo, and Cytokine-only on PBMC. **(C)** Unified training compared with training restricted to cancer, immune or H1 cellular contexts. The immune-context comparison is the same PBMC comparison shown in B. **(D)** Model comparison on three gene-editing benchmarks. GEARS [27] and X-Cell [14] are evaluated on VCC and HepG2. Evaluation configurations for PIE [13] and the other models are described in Appendix B.4. **(E)** Drug-response prediction in five held-out Tahoe-100M cell lines, covering single-drug conditions and Sacubitril-Valsartan combinations. **(F)** Cytokine-response prediction in PBMCs. D–F evaluate predictions across all measured genes, with predictions for genes outside a model’s output set filled using matched control expression. UPM achieves the highest PES among the compared models and improves all four metrics over State in E [15] and Stack in F [16]. MMD evaluates single-cell distribution prediction; ΔPearson and ΔPert *r* measure control-centered mean-response correlations across genes and conditions, respectively; PES measures perturbation discrimination. Lower MMD and higher values of the other metrics indicate better performance. Red outlines mark the best displayed scores. MMD uses logarithmic axes in A–D and linear axes in E–F.

#### Full-gene models show lower MMD than HVG2000 models across both evaluation panels, with dataset-dependent differences in mean-response correlations and perturbation discrimination

The gene-channel comparison used genetic perturbation data from Adamson, Replogle, Nadig, Norman and VCC, together with chemical perturbation data from sci-Plex 3 and SciPlexCombo [5, 6, 19, 20, 21, 23, 24]. Models were trained with supervision on 200 highly variable genes (HVG200), 2,000 highly variable genes (HVG2000), or the full reference gene set. HVG supervision restricted the genes contributing to the reconstruction loss while retaining all input and output gene channels. All three settings were evaluated on the HVG200 panel, and HVG2000 and full-gene training were also compared on the HVG2000 panel (Figure 2A; Table S1). Checkpoint selection and gene-mapping details are provided in Appendix B.4. On the HVG200 evaluation panel, expanding supervision from 200 to 2,000 genes increased MMD across all three datasets. In contrast, full-gene training reduced MMD relative to HVG2000 training across both evaluation panels, with reductions of 38.1–76.4%. Both mean-response correlations improved on VCC and HCT116 but decreased on HepG2. PES decreased in five of the six dataset–panel comparisons.

#### Joint training across perturbation types improves bulk-level metrics

This comparison used an expanded training collection that additionally included X-Atlas/Orion genetic perturbations and Parse PBMC cytokine responses [22, 26]. Unified training was compared with training restricted to the corresponding perturbation type on genetic, chemical-combination and cytokine benchmarks (Figure 2B). Both mean-response correlations improved across all three benchmarks. Chemical combinations showed improvements across all four metrics, including 14.6% higher PES and 12.3% lower MMD. For genetic and cytokine perturbations, MMD increased slightly, while PES decreased slightly on genetic perturbations and increased slightly on cytokine perturbations.

#### Joint training across cellular contexts further improves bulk-level metrics

Using the same expanded collection, unified training was compared with training restricted to cancer, immune or H1 cellular contexts (Figure 2C). Both mean-response correlations and PES improved across all evaluated cellular contexts, with the largest correlation gains on H1. MMD decreased by 67.2% on H1 but increased slightly on HepG2, HCT116 and PBMC.

#### Together, unified modeling yields a single model that achieves SOTA distributional and bulk-level performance across diverse benchmarks spanning genome-wide perturbations, perturbation types, and cellular contexts

The final unified model was trained on the expanded collection together with expression data from 45 Tahoe-100M cell lines [25] and compared with existing predictors across genetic, drug and cytokine benchmarks [13, 14, 15, 16, 27, 28, 29]. Training procedures and checkpoints for all comparison models are described in Appendix B.4. UPM achieved the highest ΔPert *r* and PES on VCC, HepG2 and HCT116, with PES exceeding the strongest displayed comparator for each dataset by 4.4–16.9% (Figure 2D). UPM also achieved the highest ΔPearson on HepG2 and HCT116, while Geneformer performed best on VCC. MMD was lower than X-Cell on VCC and HepG2, although GEARS achieved the lowest values on both datasets. UPM achieved the lowest MMD on HCT116. On the five held-out Tahoe-100M cell lines, UPM improved all four metrics over State, with ΔPearson increasing from 0.082 to 0.540 and PES improving by 56.1% (Figure 2E). On PBMC, UPM improved all four metrics over Stack, including a 37.3% gain in PES (Figure 2F). PerturbMean retained higher ΔPert *r* on Tahoe and higher values of both mean-response correlations on PBMC.

### For combinatorial perturbations, UPM achieves increasingly greater predictive advantages as the perturbation combinations become more “difficult”

#### A workflow was designed to quantify the difficulty of predicting combinatorial perturbation effects and evaluate model performance across difficulty levels

The evaluation covers simultaneous perturbation of two genes (gene-gene) and genetic perturbation combined with drug treatment (gene-drug), representing combinations within and across perturbation types, respectively. Combination difficulty is defined by the departure of observed responses from additivity (Figure 3A). The Additive baseline sums measured single-perturbation expression changes. Non-additivity is quantified as one minus the cosine similarity between the observed combination response and the sum of its constituent single-perturbation responses. Larger distances indicate more difficult combinations. Model scores and the fraction of predictions exceeding each comparator are evaluated across difficulty bins. The analysis covers 131 Norman gene-gene pairs and 16,185 genes, and 12,083 sci-Plex-GxE gene-drug tasks and 18,904 genes across A172, T98G and U87MG cells [20, 30]. Both single-gene perturbations in each Norman pair were represented in training. MMD assesses single-cell distribution prediction, while ΔPearson and ΔPert *r* assess condition-wise and gene-wise mean-response correlations. Additive and PerturbMean are represented as point distributions for MMD. For gene-drug combinations, an additional baseline, Gene-only + Drug-only, sums the expression changes independently predicted by Gene-editing-only and Molecule-only and adds the combined change to the matched unperturbed cell state. Full evaluation details are provided in Appendix B.4.1.

**Figure 3:**
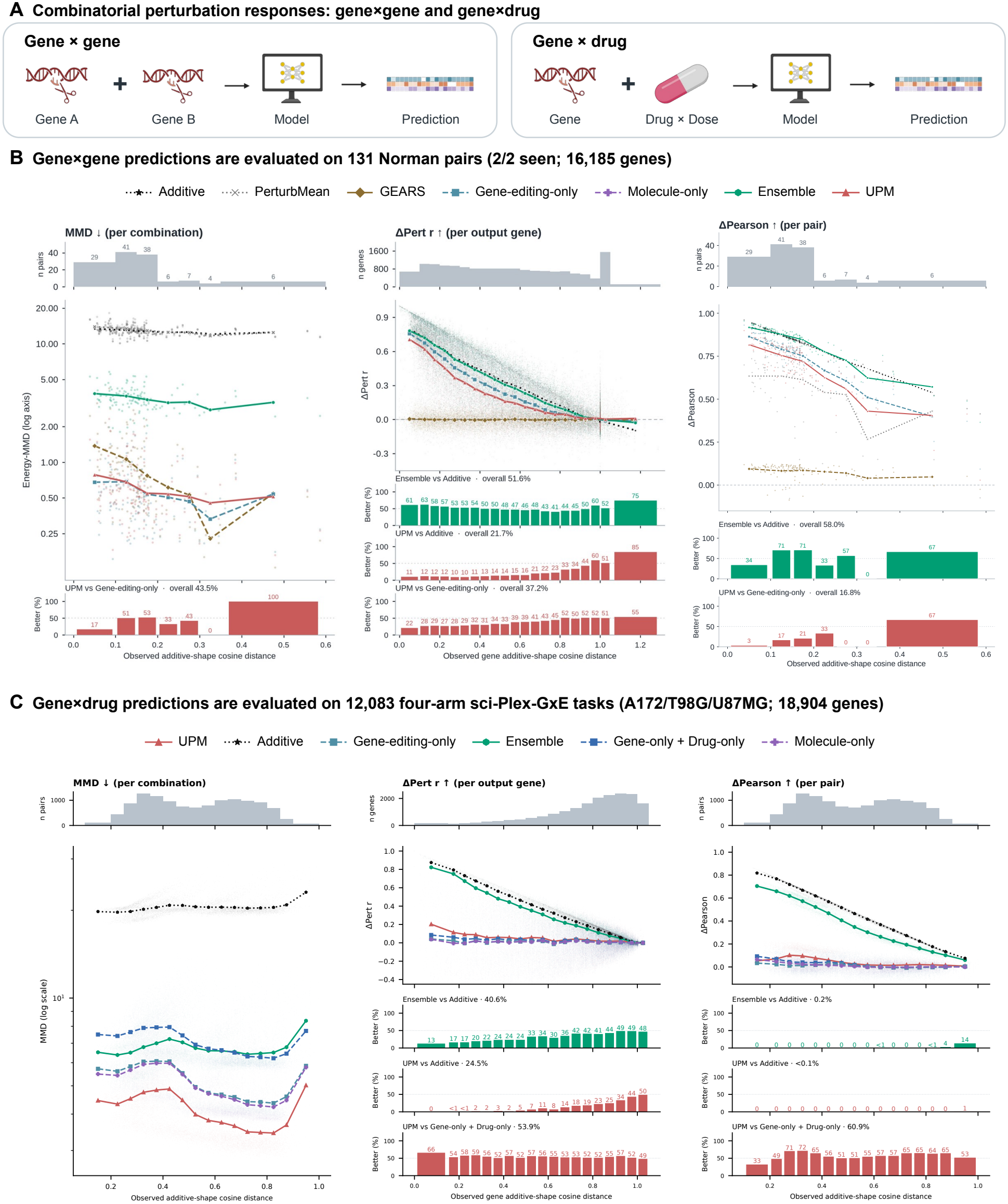
The fraction of genes better predicted by UPM than by Additive increases with combination non-additivity. **(A)** Prediction settings for combinations of two genetic perturbations or a genetic perturbation with a drug treatment. **(B)** Evaluation of 131 Norman gene-gene pairs across 16,185 genes. Both constituent single-gene perturbations are represented in training. **(C)** Evaluation of 12,083 sci-Plex-GxE gene-drug tasks across 18,904 genes in A172, T98G and U87MG cells. Combination difficulty is quantified by non-additivity, defined as one minus the cosine similarity between the observed combination response and the sum of its constituent single-perturbation responses. Distances are calculated across genes for condition-wise analyses and across conditions for gene-wise analyses; larger values indicate greater non-additivity. (Caption continued on the next page *↓*.) (Caption continued from the previous page *↑*.) Histograms show the number of observations in each difficulty bin, with individual scores shown as points and bin means as lines. Metrics are defined in Figure 2. MMD is plotted on a logarithmic scale. Lower bars report the percentage of predictions outperforming each named comparator within bins, alongside the overall percentage. Comparisons with 0% or 100% wins in every bin are omitted from these bars. Additive sums measured single-perturbation expression changes. In C, Gene-only + Drug-only sums changes predicted independently by Gene-editing-only and Molecule-only before adding them to the matched unperturbed cell state; both individual models are also shown. Additive and PerturbMean are represented as point distributions for MMD calculation. Ensemble combines Additive with two UPM configurations using weights of 50%, 25% and 25%, respectively.

#### The additive baseline performs strongly on easy perturbation combinations, while its predictions become unreliable as the difficulty increases

Additive achieved the highest overall mean-response correlations among standalone predictors in both gene-gene and gene-drug combinations. Its gene-wise correlations nevertheless decreased as observed responses departed further from additivity, and a growing fraction of output genes favored UPM in ΔPert *r*. In gene-gene combinations, this fraction reached 85% in the highest displayed gene-wise difficulty bin (Figure 3B). In gene-drug combinations, it reached 49.8% in the highest displayed bin (Figure 3C). UPM achieved lower MMD than Additive in both gene-gene and gene-drug combinations.

#### Ensembling additive approach and model predictions rescues predictive performance for difficult perturbation combinations

The Ensemble combines measured Additive responses with two UPM configurations at weights of 50%, 25% and 25%. Ensemble improved gene-wise correlations over Additive for subsets of output genes in both combination settings. Improvements in condition-wise correlation were more widespread for gene-gene combinations, occurring in 58.0% of pairs, compared with 0.2% of gene-drug tasks (Figure 3B–C). For gene-drug combinations, this fraction reached 14.1% in the highest displayed difficulty bin.

#### In gene–gene combinatorial perturbation prediction, unified modeling outperforms specialized genetic perturbation predictors, with its predictive advantage increasing as perturbation combination difficulty increases

Compared with Gene-editing-only, the fraction of pairs with lower MMD under UPM increased from 17% in the lowest displayed difficulty bin to 100% in the highest (Figure 3B). The fractions of output genes and perturbation pairs favoring UPM in ΔPert *r* and ΔPearson, respectively, were also higher in the highest than in the lowest displayed difficulty bins, indicating greater benefits for strongly non-additive responses.

#### In gene-drug combinatorial perturbation prediction, unified modeling outperforms the combined predictions of specialized genetic and drug perturbation models overall, and its advantage over Additive increases with combination difficulty

Gene-only + Drug-only combines independently predicted genetic and drug effects, while Additive uses measured single-perturbation responses. Relative to Gene-only + Drug-only, UPM reduced mean MMD by 41.7% and achieved lower MMD in all 12,083 tasks (Figure 3C). Compared with Gene-only + Drug-only, UPM achieved higher ΔPert *r* for 53.9% of output genes and higher ΔPearson for 60.9% of tasks. With increasing difficulty, the fraction of genes favoring UPM in ΔPert *r* increased relative to Additive but generally decreased relative to Gene-only + Drug-only. The ΔPearson win fraction against Gene-only + Drug-only did not increase consistently with difficulty.

### UPM recapitulates HSC perturbation-response directions with 40.0% higher pooled accuracy than Gene-editing-only

#### Simulations are performed to computationally examine the responses of human HSCs to gene knockdown and cytokine stimulation, and compare the predicted responses with experimental results from humans and mice

Both simulation settings start from human hepatic stellate cell (HSC) expression states associated with metabolic dysfunction-associated steatohepatitis (MASH; Figure 4A). The first combines GAS7, SERPINE1, SPON1 or LTBP2 knockdown with TGF*β*1 stimulation and evaluates ACTA2, COL1A1, SERPINE1 and LOXL2 responses against published human HSC experiments [17]. The second simulates ETS1, ETS2, GATA4 or GATA6 knockdown and evaluates COL1A1, LOXL2, IL6 and CXCL5 responses against published mouse HSC experiments [18]. The latter compares predictions from human HSCs with response directions measured in mouse HSCs. Each setting contributes 16 target-readout pairs, scored relative to random-knockdown references. In the combined setting, Gene-editing-only predicts the knockdown response from the same HSC input, while UPM receives both the knockdown and TGF*β*1 stimulation. X-Cell [14] is compared in the single-KD setting.

**Figure 4:**
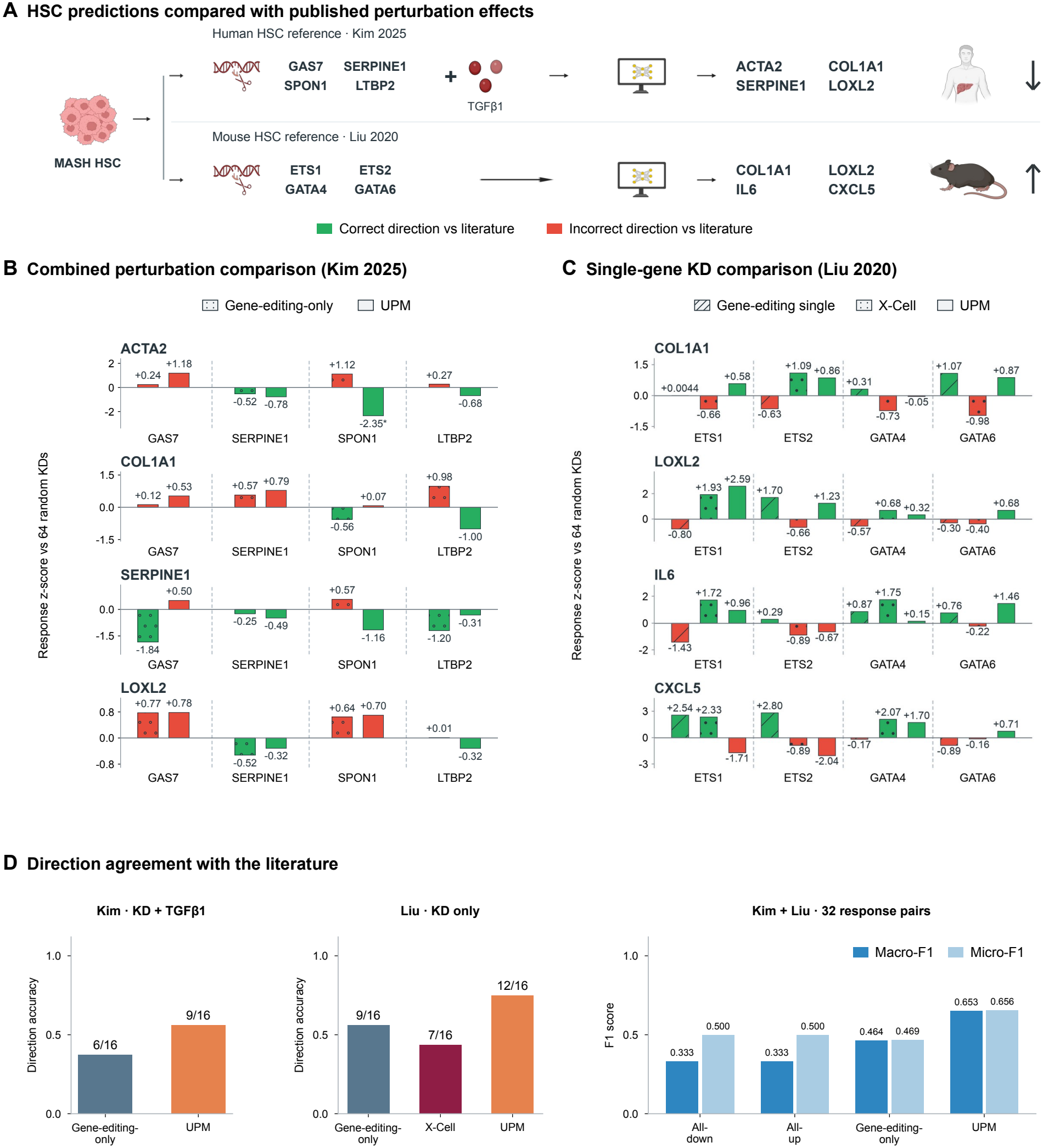
UPM recovers HSC perturbation-response directions with higher pooled accuracy than Gene-editing-only. **(A)** Both simulation settings start from human MASH hepatic stellate cell (HSC) expression states. Predictions for combined gene knockdown and TGF*β*1 stimulation are compared with human HSC experiments [17]; predictions for single-gene knockdowns are compared with mouse HSC experiments [18]. **(B)** Predicted ACTA2, COL1A1, SERPINE1 and LOXL2 responses to GAS7, SERPINE1, SPON1 or LTBP2 knockdown. Gene-editing-only models the knockdown alone, and UPM models the knockdown together with TGF*β*1 stimulation. **(C)** Predicted COL1A1, LOXL2, IL6 and CXCL5 responses to ETS1, ETS2, GATA4 or GATA6 knockdown, comparing Gene-editing-only, X-Cell [14] and UPM. Bars in B–C show response z-scores relative to 64 random-knockdown controls. Green indicates agreement with experimentally reported response directions, and red indicates disagreement. Bar patterns distinguish models. **(D)** Directional accuracy is reported separately for combined interventions evaluated against human experiments and single-gene knockdowns evaluated against mouse experiments. Pooled macro-F1 and micro-F1 (equal to directional accuracy) are calculated across all 32 target-readout pairs, with all-down and all-up predictors as baselines. UPM correctly predicts 21/32 response directions compared with 15/32 for Gene-editing-only, a 40.0% relative increase in accuracy; their macro-F1 scores are 0.653 and 0.464, respectively.

#### Virtual cell simulations recapitulate the directions of 21 out of 32 perturbation responses reported in human and mouse HSC studies

In the combined knockdown and TGF*β*1 stimulation setting, UPM recovered 9/16 reported response directions, compared with 6/16 for Gene-editing-only (Figure 4B). These included reductions in ACTA2, COL1A1 and SERPINE1 following SPON1 knockdown. In the single-KD setting evaluated against the mouse HSC study, UPM recovered 12/16 directions, compared with 9/16 for Gene-editing-only and 7/16 for X-Cell (Figure 4C). GATA6 knockdown matched the published directions for all four readouts.

#### UPM achieves the highest pooled directional accuracy and macro-F1 among the predictors evaluated across both HSC settings

Across 32 target-readout pairs, UPM achieved 65.6% accuracy (21/32), compared with 46.9% (15/32) for Gene-editing-only, representing a 40.0% relative improvement (Figure 4D). Macro-F1 was also higher for UPM (0.653 versus 0.464) (Table S2). Constant all-up and all-down predictions each achieved 50.0% accuracy and macro-F1 of 0.333. Evaluation details are provided in Appendix B.4.2.

## Discussion

This study presents a unified modeling approach that integrates heterogeneous perturbation resources to broaden AIVC predictive capabilities. In the training-scope comparisons, full-gene supervision improved single-cell distribution prediction, while broader coverage of perturbation types and cellular contexts improved mean-response prediction. Expanding cellular context coverage also improved perturbation discrimination. The unified model achieved strong performance across genetic, chemical and cytokine perturbation benchmarks. Together, these findings demonstrate the effectiveness of UPM in improving AIVC prediction across gene sets, perturbation types and cellular contexts.

UPM extends unified modeling to perturbation combinations within and across perturbation types. Its advantage over Gene-editing-only was more pronounced for highly non-additive gene-gene responses. For gene-drug combinations, joint modeling improved both distribution prediction and overall mean-response correlations relative to summing independently predicted genetic and drug effects. Across both settings, a growing fraction of genes favored UPM over Additive as non-additivity increased. Further improvements from combining UPM with Additive in gene-gene prediction illustrate the complementarity between learned combination responses and measured single-perturbation effects.

The HSC application extends this analysis to genetic perturbations with and without cytokine stimulation in a disease-associated setting. Starting from human MASH HSC expression states, UPM recovered more published response directions than Gene-editing-only in both single-gene knockdown and combined knockdown and cytokine stimulation settings, using human and mouse studies as experimental references. UPM provides an in silico framework for simulating candidate therapeutic interventions in disease-associated cellular contexts, with potential applications in drug target prioritization and the design of combination therapies.

Despite its advantages, UPM and its evaluation have several limitations. Several external models were evaluated using publicly released checkpoints trained on datasets different from those used for UPM. Some models’ output gene sets also did not cover all genes included in the evaluation, so predictions for uncovered genes were filled with the corresponding control expression values. Comparisons with a broader range of VAE-based models and LLM-based perturbation predictors are still lacking. For combinatorial perturbation prediction, Additive retains the highest overall mean-response correlations among standalone predictors. Predicted responses are not always sufficiently specific to individual perturbations, a pattern also reflected in the UMAP projections. The evaluation remains limited in its range of metrics. A broader assessment could incorporate the calibrated metrics discussed by Miller et al. [31] and the Virtual Cell Challenge 2026 metric suite [32] to better characterize perturbation-specific responses and differential-expression accuracy.

## Acknowledgements

This work was supported in part by the University Development Fund–Research Start-up Fund of The Chinese University of Hong Kong, Shenzhen (Grant No. UDF01004259).

## Appendix

### A Related work

Perturbation predictors differ in their representations of cell states and interventions. scGen combines a VAE with latent vector arithmetic across cellular contexts [33]; CPA composes perturbation, dose and covariate representations for unseen combinations [24]. chemCPA predicts single-cell responses to unseen drugs [34]. GEARS and PIE use biological relationships or priors [13, 27]. Geneformer and scGPT learn transferable expression representations [28, 29], and State and Stack use large-scale training or contextual information [15, 16].

Existing approaches already address multiple aspects of unified perturbation modeling. X-Pert and UniPert-G2CP connect genetic and chemical perturbations through shared representations [10, 35]; PRnet models chemical responses across cellular contexts [36], while CellFlow and biolord provide frameworks applicable to multiple perturbation settings [12, 37]. Against this background, UPM examines the empirical benefits of jointly modeling heterogeneous gene sets, perturbation types and cellular contexts within a shared architecture.

Protein language models complement expression representations with sequence information. ESM-2 learns protein-sequence embeddings [38]. UPM uses fixed sequence features with a trainable projection and shared perturbation encoder for genetic and cytokine inputs.

Generative models predict changes in cellular distributions. CellOT learns neural transport maps between unpaired populations [39]; CellFlow uses conditional flow matching [37], and scDFM combines flow matching with MMD [40]. Squidiff uses diffusion-based denoising [41], while X-Cell and Lingshu-Cell model perturbed expression states generatively [14, 42]. Virtual tissue models extend this scope to spatial organization: VirTues learns multiscale spatial-proteomic representations [43], and CIFM models cell-cell interactions under combinatorial perturbations [44]. These approaches motivate evaluating cellular distributions alongside mean responses.

Metrics and baselines affect conclusions about predictive progress. Linear and control predictors can match or exceed deep models in some genetic benchmarks [45, 46]. Systema separates perturbation-specific responses from shared systematic variation [47]; calibrated metrics can reveal deep-model gains over uninformative baselines [31]. The present evaluation combines MMD for distributions, PES for perturbation discrimination, and ΔPearson and ΔPert *r* for condition-wise and gene-wise correlations. PerturbMean and Additive are included where applicable, with combinations evaluated across non-additivity levels.

Benchmarks also differ in generalization scope. Virtual Cell Challenge 2025 holds out genetic perturbations within one cellular context [21]; the 2026 challenge targets zero-shot prediction across contexts [48]. Each comparison therefore specifies training data, perturbation inputs, output genes and holdouts to distinguish new-perturbation from new-context generalization.

### B Method

#### B.1 Unified perturbational modeling and model architecture

**Figure S1:**
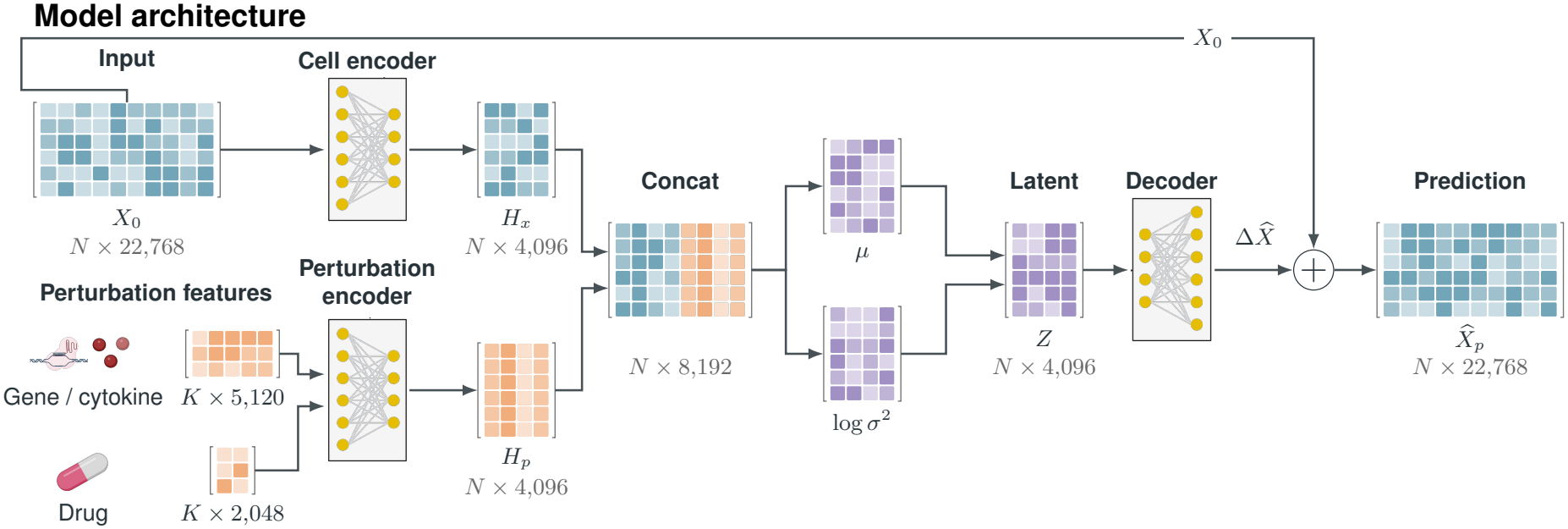
UPM model architecture and tensor dimensions. A cell-state encoder maps unperturbed expression profiles to 4,096-dimensional representations. Genetic and cytokine inputs use 5,120-dimensional protein features, and drugs use 2,048-dimensional Morgan fingerprints. Projected features pass through a shared perturbation encoder. Constituent embeddings are scaled, averaged and concatenated with each cell-state representation to parameterize the mean and log variance of a diagonal Gaussian through separate networks. Training samples this distribution; inference uses its mean. The decoder predicts expression changes that are added to the input expression. Core networks use six-layer (L6) or ten-layer (L10) architectures. *N* denotes the number of input cells for one perturbation condition, and *K* denotes the number of constituents of the corresponding feature type.

##### Gene alignment and masking

UPM aligns expression profiles across datasets to a shared gene space constructed from the union of measured genes, matched by their Ensembl identifiers. Each gene occupies the same position in the aligned expression vectors across datasets. The final unified model represents 22,768 genes, and each benchmark is evaluated on its specified set of measured genes.

For dataset *d*, let *G*_*d*_ denote its measured gene set and =*U*_*d*_ *G*_*d*_ the shared gene set. The mapping *π*_*d*_(*g*) assigns each measured gene to its position in the aligned expression vector, with zeros inserted for genes not measured in that dataset. A source-specific Boolean mask restricts training losses to mapped genes with positive total expression across the source. Zero expression in an individual cell remains a supervised observation if the gene is retained by this mask. The mask is applied only when computing the losses and is not concatenated to the encoder input. For HVG supervision, the retained gene set is further restricted to the fixed training HVG list.

##### Cell-state encoding

The shared cell-state encoder maps an unperturbed expression profile *x*_0_ ∈ ℝ^*G*^ in the shared gene space to a latent representation *h*_*x*_ = *f*_*x*_(*x*_0_). In its first linear layer,

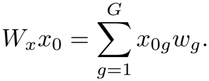

Here *w*_*g*_ is the trainable weight vector associated with gene *g*. Each gene therefore contributes through its expression value and a gene-specific weight vector, without a separate gene-identity embedding network. Protein-sequence features encode genetic perturbations through a separate input pathway, as described below.

##### Network architecture

UPM comprises five residual MLPs: a cell-state encoder, a perturbation encoder, separate networks for the latent mean and log variance, and an expression decoder (Figure S1). Each MLP contains six linear layers in L6 or ten in L10, with a hidden width of 4,096, ReLU activations and LayerNorm. The cell-state encoder maps *G*-gene expression profiles to 4,096-dimensional representations. Two-layer feature projections map perturbation features to 4,096 dimensions before the shared perturbation encoder. The cell-state and perturbation representations are concatenated into an 8,192-dimensional input to the mean and log-variance networks, each producing 4,096-dimensional outputs. The decoder maps the resulting 4,096-dimensional latent representation to expression changes across *G* genes.

##### Genetic and cytokine features

Genetic perturbations use fixed 5,120-dimensional protein-sequence embeddings from ESM-2 (esm2_t48_15B_UR50D; https://huggingface.co/facebook/esm2_t48_15B_UR50D) [38]. Each gene is mapped to a set of UniProt proteins U(*g*), whose embeddings are averaged:

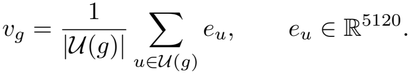

Cytokines use precomputed 5,120-dimensional features. Both feature types remain fixed during training and pass through a shared trainable two-layer projection to 4,096 dimensions, followed by the shared perturbation encoder.

##### Chemical features and dose encoding

Chemical perturbations use fixed 2,048-bit Morgan fingerprints generated from SMILES using RDKit, with radius 3 and chirality excluded [49] (RDKit documentation). A trainable two-layer projection maps the fingerprints to 4,096 dimensions before the shared perturbation encoder. The encoded representation is scaled by a dose coefficient:

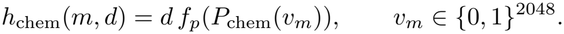

The coefficient is *d* = ln(1 + *c/*100), where *c* is the drug concentration in nM. Each drug in a combination is scaled by its own dose before aggregation.

##### Perturbation encoding and aggregation

Each constituent perturbation *p*_*k*_ is encoded and scaled before aggregation:

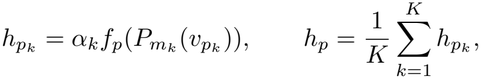

where 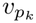 is the input feature, 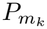 is the corresponding feature projection, and *f*_*p*_ is the shared perturbation encoder. The coefficient *α*_*k*_ is −1 for CRISPRi, 1 for CRISPRa and cytokines, and *d*_*k*_ = ln(1 + *c*_*k*_*/*100) for drugs, with concentration *c*_*k*_ in nM. Averaging the *K* adjusted embeddings yields a single representation for perturbations within or across types, with *K* = 1 for a single perturbation (Figure 3).

##### Latent fusion and expression decoding

Separate networks map the concatenated representations [*h*_*x*_, *h*_*p*_] to the mean *µ* and log variance *ℓ* of a diagonal Gaussian:

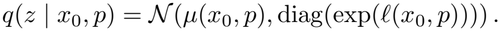

Training samples *z* = *µ* + exp(*ℓ/*2) ⊙ *ϵ*, with *ϵ* ∼ *N*(0, *I*), and applies KL regularization against a standard-normal prior:

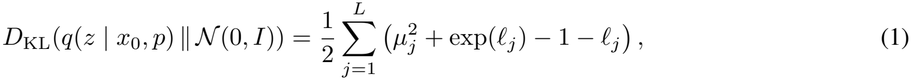

where *L* is the latent dimension. Inference uses the latent mean, and the decoder predicts an expression change that is added to each input cell:

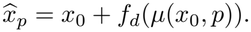

Predictions retain the input gene ordering, with population variability arising from differences among input cell states.

##### Training objective

For condition *b*, control cells *C*_*b*_ and perturbed cells *P*_*b*_ are independently sampled from the same source and cellular context. Training compares predictions *Y*_*b*_ generated from *C*_*b*_ with *P*_*b*_ through population distributions and mean responses, without cell-level pairing. Losses are restricted to the gene set *S*_*b*_ retained by the measurement and supervision masks. For cell populations 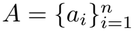 and 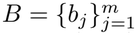, with equal weights within each population, the distribution loss is the GeomLoss energy discrepancy computed on these genes:

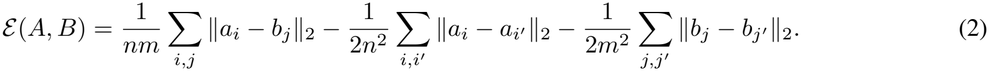

Mean-response error is measured by

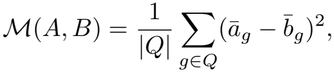

where *ā*_*g*_ and 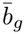 are predicted and observed mean expression, respectively. The evaluated gene set *Q* excludes genes with zero observed means and negative predicted means. For each condition *b, T*_*b*_ ⊆ *S*_*b*_ selects genes with the largest absolute observed changes 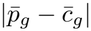, retaining the top 10% or at least 1,000 genes, capped by the available gene count:

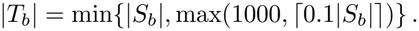

For the expression-change models, *Y*_*b*_ = *C*_*b*_ + *f*_*d*_(*z*_*b*_) and the complete objective is

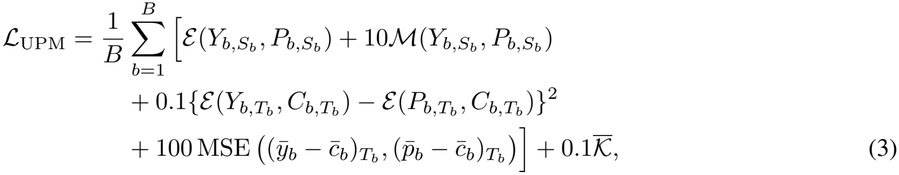

where *B* is the number of conditions in the device batch and MSE averages over genes. The normalized latent penalty averages over all *N* input control cells and all *L* = 4096 latent dimensions:

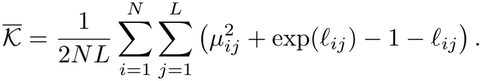

The four response-loss weights are 1, 10, 0.1 and 100, and the normalized KL weight is 0.1. All weights are constant throughout training. Distribution losses use the raw decoded values; nonnegative clipping is applied when

##### Gene-supervision objective

The three models in Figure 2A decode absolute expression. Let *R*_*b*_ and *Y*_*b*_ denote control reconstructions at zero perturbation and perturbed reconstructions, respectively. Their objective follows Equation 3, but uses *R*_*b*_ instead of *C*_*b*_ as the predicted reference in 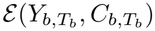 and 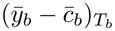, and adds the control reconstruction loss

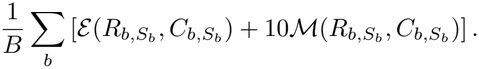

The observed reference remains *C*_*b*_. Training multiplies decoded reconstructions by gene-wise Bernoulli masks. For control mean *u*_*g*_ and mean positive control expression *v*_*g*_, 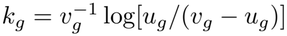, with invalid coefficients set to ™∞. Retention probabilities are *σ*(*k*_*g*_*v*_*g*_) for controls and 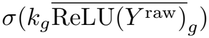 for perturbed cells. The latter mean is detached from differentiation, and non-finite probabilities are set to zero. Control reconstruction uses the latent mean; perturbed reconstruction uses a sampled latent state. Evaluation uses latent means and unmasked outputs:

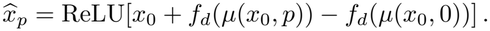

Figure 2A evaluates these predictions without additional library-size renormalization.

#### B.2 Perturbation datasets, preprocessing and data splits

##### Collection size and exclusions

The resource contains 110,069,606 expression profiles across 64 files. Held-out data, controls with no retained training conditions and non-control conditions with fewer than ten cells are excluded from training. After preprocessing and splitting, the training set contains 89,652,569 profiles: 86,834,887 perturbed cells and 2,817,682 matched controls.

##### Gene and cellular-context coverage

The shared gene space contains 22,768 unique Ensembl genes. Training covers 71 cellular contexts (53 cell lines and 18 PBMC cell types), and the complete resource covers 76 (58 cell lines and the same 18 cell types). Contexts are deduplicated across datasets independently of donor identity. Figure 1C shows the resource total and within-dataset counts. PBMC donors are counted separately: eight in training and twelve in the complete resource.

##### Training datasets

Genetic training data include Adamson K562 [5], Replogle K562/RPE1 [6], Nadig HepG2/Jurkat [19], Norman K562 [20], Virtual Cell Challenge 2025 H1 [21], and X-Atlas/Orion HCT116/HEK293T [22]. Chemical training data include sci-Plex 3 A549/MCF7/K562 [23], retained SciPlexCombo individual-drug conditions in A549 [24], and 45 Tahoe-100M cell lines [25]. Cytokine data come from Parse PBMC [26]. sci-Plex-GxE is reserved for gene-drug evaluation [30].

Figure 2A trains on Adamson K562 [5], Replogle K562/RPE1 [6], Nadig HepG2/Jurkat [19], Norman single-gene conditions [20], VCC H1 training conditions [21], sci-Plex 3 A549/MCF7/K562 [23], and SciPlexCombo A549 individual-drug conditions [24]. This collection excludes X-Atlas/Orion, Parse PBMC and Tahoe expression profiles.

Panels B and C add X-Atlas/Orion HCT116/HEK293T [22] and Parse PBMC [26], but exclude Tahoe. Gene-editing-only retains the genetic sources; Molecule-only retains sci-Plex 3 and SciPlexCombo individual-drug conditions; Cytokine-only and Immune-only share the Parse PBMC subset. Cancer-only retains A549, HCT116, HepG2, Jurkat, K562 and MCF7 from Adamson, Replogle, Norman, Nadig, X-Atlas/Orion, sci-Plex 3 and SciPlexCombo. H1-only uses VCC H1 training data. All subsets inherit their parent collection’s condition exclusions.

Panels D–F add the 45 Tahoe-100M training cell lines listed below [25], giving ten data resources. All three external benchmarks use the same final checkpoint.

##### Cell lines, cell types and donors

The datasets other than Tahoe cover nine distinct cell lines: A549, H1, HCT116, HEK293T, HepG2, Jurkat, K562, MCF7, and RPE1. The 45 Tahoe training cell lines are A172, A427, A498, A549, AN3CA, AsPC1, BT474, C33A, CFPAC1, CHP212, COLO205, H4, HCT15, HEC1A, HS578T, HT29, hTERTHPNE, J82, KATOIII, LoVo, LOXIMVI, LS180, MIAPaCa2, NCIH1573, NCIH1792, NCIH2030, NCIH2122, NCIH23, NCIH2347, NCIH460, NCIH596, NCIH661, Panc03.27, RKO, RPMI7951, SHP77, SKMEL2, SNU1, SNU423, SW1088, SW1271, SW1417, SW48, SW480, and SW900.

PBMC contains 18 cell types: B Naive, B Intermediate/Memory, CD4 Naive, CD4 Memory, CD8 Naive, CD8 Memory, Treg, MAIT, NKT, NK, NK CD56bright, CD14 Mono, CD16 Mono, cDC, pDC, HSPC, ILC and Plasmablast. Training uses Donors 1, 3, 5, 6, 7, 9, 11 and 12. Filtering conditions with fewer than ten cells leaves no training conditions for ILC-Donor6, Plasmablast-Donor5, Plasmablast-Donor9, Plasmablast-Donor11 or pDC-Donor7.

##### Perturbation conditions

Training contains 83,484 non-control conditions defined within each source and cellular context. Genetic perturbations contribute 19,542 conditions and 2,769,801 cells, covering 11,730 target-gene labels or 11,783 identities when CRISPRi and CRISPRa are distinguished; no gene-gene combinations are included. Individual drugs contribute 52,698 conditions and 77,821,532 cells across 1,853 drug-dose labels and 551 drug names. Sacubitril plus Valsartan contributes 134 conditions and 203,027 cells, with both components assigned doses of 50, 500 or 5,000 in the source metadata. Cytokines contribute 11,110 conditions and 6,040,527 cells across 90 cytokines. Matched control pools contain 475,904 genetic, 1,912,832 chemical and 428,946 PBMC cells.

##### Validation and test splits

Panel A reserves predefined HepG2 targets for validation and testing [19] and excludes all Norman gene-gene and SciPlexCombo drug-drug combinations from expression training [20, 24]. For SRT3025, PCI-34051, Sorafenib, Dasatinib, SRT2104, SRT1720, Crizotinib, Alvespimycin and Curcumin, dose labels 10 and 100 are used for validation, and 1,000 and 10,000 for testing. VCC validation non-control conditions and test expression profiles are held out; test gene identifiers contribute to the shared gene set [21]. HCT116 is an unseen cellular context because this collection excludes X-Atlas/Orion expression data [22].

Panels B and C retain the panel A splits and add held-out HCT116 targets [22] and PBMC donor splits [26]. PBMC Donors 1, 3, 5, 6, 7, 9, 11 and 12 are used for training, 4 and 10 for validation, and 2 and 8 for testing. Restricted models retain these splits within their selected sources. Tahoe expression profiles are excluded.

Panels D–F retain the panel B/C splits and add 45 Tahoe-100M training cell lines. C32, HOP62, HepG2-C3A, Hs 766T and PANC-1 are held out [25]. The Tahoe Sacubitril-Valsartan combination remains in training. Perturbation holdouts are defined within each dataset and cellular context.

##### Evaluation sets and gene panels

The broad genetic summary contains 928 conditions: 93 VCC, 476 HepG2, 228 HCT116 and 131 Norman. The perturbation-type comparisons evaluate 25 SciPlexCombo combinations on 15,763 genes and average 36 PBMC cell-type/donor units. Gene-gene evaluation covers 131 pairs and 16,185 genes, including 16,173 with finite gene-wise correlations. Gene-drug evaluation covers 12,083 conditions and 18,904 genes across 510 targets, three cell lines, four drugs and two doses. Evaluation units are defined separately for each benchmark.

External genetic benchmarks use matched subsets of 29 VCC/H1, 258 HepG2 and 207 HCT116 perturbations on 17,540, 9,191 and 18,907 genes, respectively. Tahoe evaluates five held-out cell lines on 19,668 genes; PBMC evaluates 36 donor/cell-type groups on 18,481 genes. Fixed-model HVG200 and HVG2000 analyses retain the genetic test perturbations and change only the training-derived evaluation gene panel (Appendix B.4).

#### B.3 Evaluation metrics and aggregation

##### Control-centered responses

Within each evaluation unit, let *N* denote perturbation conditions, *G* output genes, and 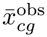 and 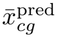 the observed and predicted population means for condition *c* and gene *g*. Control-centered responses are

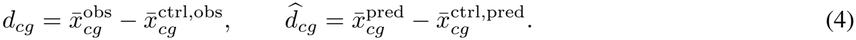

Means are computed from unpaired observed and predicted cells in the evaluation expression space. Comparisons with a shared observed control use the same control mean for both.

##### Maximum mean discrepancy (MMD)

For predicted cells 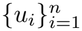 and observed cells 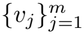, MMD uses the GeomLoss energy kernel *k*(*u, v*) = ™∥*u* ™ *v*∥_2_:

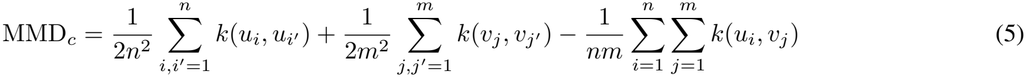

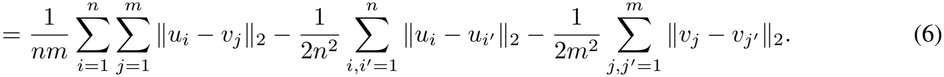

The sums include diagonal terms. The reported MMD [50] is half the squared kernel discrepancy, equivalently half the empirical energy distance, without a square root. Scores are averaged over conditions:

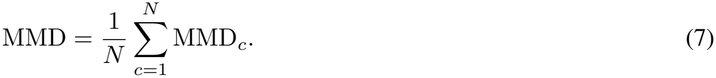

Lower MMD indicates closer distributions in both population means and cell-to-cell variation; point predictions have zero within-population distance. Native gene-drug evaluation uses matched deterministic samples capped at 32 observed and unperturbed cells. Expression space and sampling are matched within each benchmark.

##### Perturbation effect score (PES)

Let *D*_*cj*_ be the Euclidean distance from predicted response *c* to observed response *j*,

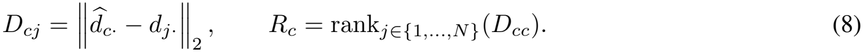

Within each row, distances are ranked from smallest to largest, with rank 1 for the closest response and average ranks for ties. Condition and mean scores are

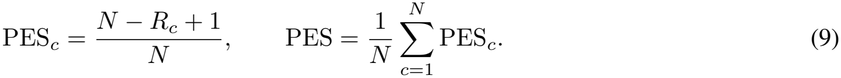

Higher PES indicates better perturbation discrimination: a unique correct nearest response scores 1, a last-ranked response 1*/N*, and a complete tie (*N* + 1)*/*(2*N* ).

##### Condition-wise ΔPearson

Define 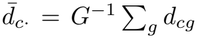 and 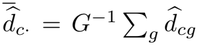. The correlation across genes for condition *c* is

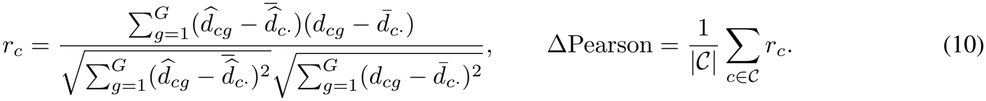

Here *C* contains conditions with valid scores under the zero-variance convention below. The metric measures agreement in response patterns across genes within each perturbation.

##### Gene-wise Δ**Pert** *r*

Define 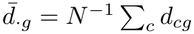 and 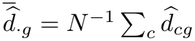. The correlation across conditions for gene *g*

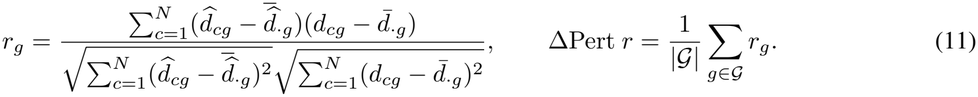

Here *G* contains genes with valid scores. The metric measures agreement in each gene’s response across perturbations. Both correlations exclude constant observed vectors and assign zero to constant predictions when observations are nonconstant. Thus, PerturbMean has ΔPert *r* = 0 in the one-context Norman task. These rules handle zero denominators; higher scores are better.

##### Score aggregation and relative gains

Genetic scores use condition-count weights across benchmarks; PBMC perturbation-type scores use equal weights across 36 units. For positive reference scores, relative increases are 100(*s*_UPM_*/s*_ref_ ™ 1)%, and relative MMD reductions are 100(1 ™ *s*_UPM_*/s*_ref_ )%. Accuracy differences in percentage points are reported separately. Win fractions count eligible genes or conditions with strictly better predictions than the reference.

For evaluation units *u* = 1, …, *U* with scores *s*_*u*_, the reported weighted mean is

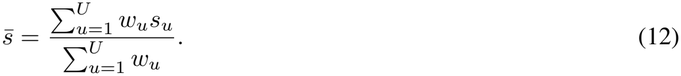

Condition-count weighting sets *w*_*u*_ = *N*_*u*_, the number of evaluated conditions; equal PBMC-group weighting sets *w*_*u*_ = 1. Correlations are computed within each evaluation unit before aggregation.

#### B.4 Training settings and evaluation benchmarks

##### Optimization and batch sampling

All UPM configurations use Adam with learning rate 10^*™*6^, *β*_1_ = 0.9, *β*_2_ = 0.999, *ϵ* = 10^*™*8^ and zero weight decay. Training uses mixed float16 precision, gradient-norm clipping at 5 and one update per batch, with fixed learning rate and loss weights. Each device processes 16 conditions. Final-collection L6/L10 and Figure 2B–C Unified models use four devices (64 conditions per update); Figure 2A and restricted models use two (32 conditions).

Each batch samples source/cellular-context entries uniformly with replacement, then conditions uniformly within each entry. Conditions are sampled without replacement when the requested count is smaller than the pool, and with replacement otherwise. Each condition independently supplies up to 128 control and perturbed cells, giving at most 2,048 of each per device batch. Incomplete final chunks of control and perturbed cells are discarded during training-data preparation. The condition-index list defines epoch length, with balanced resampling each epoch. Validation traverses held-out conditions and samples up to 1,024 control and perturbed cells each.

##### Checkpoint selection

After each epoch, validation computes Spearman correlation between observed mean expression and mean nonnegative predictions over eligible genes. Correlations are averaged within batches, then across the epoch. The highest score determines the training-selected checkpoint; the last epoch is saved separately.

Figure 2A uses the validation-selected HVG200 and HVG2000 checkpoints at completed epochs 912 and 381, respectively, and the final-epoch full-gene checkpoint at completed epoch 2,000 (502,000 optimizer updates).

The standalone models in Figure 2D–F and Figures 3–4 use the training-selected L10 checkpoint. The UPM ensemble in Figure 1D and Additive-assisted Ensemble in Figure 3 use training-selected L6 and L10 checkpoints. Gene-editing-only in Figures 3–4 uses the last-epoch genetic checkpoint from Figure 2B. Molecule-only in Figure 3 uses the validation-selected epoch 1,128 checkpoint (76,704 updates) for both standalone predictions and Gene-only + Drug-only.

For the genetic comparison in Figure 2B and Cancer-only/H1-only in panel C, training-selected and last checkpoints are compared on matched validation conditions. Each dataset–metric pair votes for lower MMD or higher PES, ΔPearson or ΔPert *r*; differences within 10^*™*9^ are ties. The checkpoint with more votes is retained. Ties are resolved by wins in ΔPearson, ΔPert *r*, MMD and PES, in that order, then by choosing the training-selected checkpoint. Validation uses VCC/HepG2/HCT116 for the genetic comparison, HepG2/HCT116 for Cancer-only and VCC for H1-only. Chemical, cytokine and immune comparisons use validation Spearman scores. Training sources and splits are given in Appendix B.2.

Completed L6 and L10 runs on the final collection contain 1,305,000 and 2,610,000 updates. Each Figure 2A run contains 502,000. Gene-editing-only, Molecule-only, Cytokine-only/Immune-only, Unified, Cancer-only and H1-only contain 1,222,000, 136,000, 696,000, 1,026,000, 520,992 and 10,000 updates, respectively.

##### Gene-supervision comparisons

Figure 2A compares supervision on 200 HVGs, 2,000 HVGs and the full gene set. HVGs are ranked by training expression variance within a shared set of 21,366 genes. Both HVG models retain all 21,366 input and output genes, but supervise only

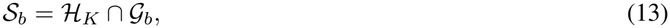

where *ℋ*_*K*_ is the fixed HVG list, *K* ∈ {200, 2000}, and *G*_*b*_ contains genes measured in batch *b*. Only *S*_*b*_ contributes to expression reconstruction loss.

The full-gene reference uses 22,768 input/output genes and one final-epoch checkpoint across all three datasets and both HVG panels. Full-gene and HVG models retain their respective training collections and gene mappings.

Evaluation maps each fixed HVG list to the dataset, excludes missing genes and ambiguous mappings, and scores all models on the same retained genes within each panel (Table S1):

**Table S1:** Numbers of genes retained for the HVG200 and HVG2000 evaluations in Figure 2A after gene mapping.

| Dataset | HVG200 | HVG2000 |
|---|---|---|
| VCC | 141 | 1,712 |
| HepG2 | 137 | 1,643 |
| HCT116 | 167 | 1,807 |

HVG200 evaluation compares all three supervision settings; HVG2000 evaluation compares HVG2000 and full-gene supervision. The fixed-model analyses below vary only evaluation genes, keeping checkpoints unchanged.

##### Perturbation-type comparisons

Figure 2B compares Gene-editing-only, Molecule-only and Cytokine-only with Unified models trained across perturbation types. All share the VAE-based architecture and use the checkpoint-selection procedures above.

##### Cellular-context comparisons

Figure 2C compares Cancer-only and H1-only with Unified models trained across cellular contexts. The immune comparison reuses the Cytokine-only models and PBMC evaluations from panel B.

##### External benchmarks

Figure 2D–F use one standalone L10 checkpoint trained on the final collection. Genetic evaluation covers matched sets of 29 VCC, 258 HepG2 and 207 HCT116 targets on all 17,540, 9,191 and 18,907 measured genes, respectively. Missing model outputs receive zero expression change. GEARS [27] and X-Cell [14] are evaluated on VCC and HepG2.

Tahoe evaluation covers 5,683 conditions across held-out C32, HOP62, HepG2-C3A, Hs 766T and PANC-1, including 15 Sacubitril-Valsartan conditions. Combination components are encoded and dose-scaled separately, then averaged before decoding. Scores are unweighted cell-line means over 19,668 genes in log1p(CP10K) space. Genes outside State’s output panel [15] retain matched control expression.

PBMC compares aligned Stack predictions [16] with final-collection L10 UPM, averaging scores equally across 36 donor/cell-type groups on 18,481 genes. External models differ in architecture, training data, output coverage and adaptation.

##### Genetic baseline fitting

In Figure 2D, Geneformer and scGPT-linear [28, 29] are fitted separately on VCC/H1, Nadig HepG2 and X-Atlas/Orion HCT116 training conditions [19, 21, 22]; GEARS [27] is fitted on VCC/H1 and HepG2. Before model-specific target mapping, the training pools contain 146, 1,128 and 9,160 conditions, respectively. Geneformer and scGPT-linear fit condition-mean responses relative to training controls, excluding predefined validation and test targets.

GEARS [27] is trained from scratch on single-cell profiles. VCC/H1 uses 187,139 training cells across 131 perturbations and 18,031 validation cells across 15 perturbations; HepG2 uses 59,360 training cells across 1,012 perturbations and 5,549 validation cells across 112 perturbations. Counts include controls. Both runs use 20 epochs, seed 1, training/validation batches of 32/128, Adam learning rate 10^*™*3^, zero weight decay and learning-rate halving each epoch. Minimum validation MSE on the top 20 differentially expressed genes selects epoch 1 for VCC/H1 and epoch 2 for HepG2. Checkpoints remain fixed across full-gene and HVG evaluations; HCT116 is omitted. Distributional predictions add mean changes to 256 sampled training-control cells and clip negative expression to zero. Code: https://github.com/snap-stanford/GEARS.

Geneformer [28] uses frozen 768-dimensional embeddings from Geneformer-V2-104M/model.safetensors, pre-trained on Genecorpus-104M. Multi-output ridge regression maps target embeddings to control-centered responses, with an intercept and regularization 1.0. scGPT-linear [29] uses frozen 512-dimensional embeddings from human scGPT best_model.pt. Cell-count-weighted bilinear ridge regression relates perturbation-gene and output-gene embeddings to responses, with regularization 0.1 and no intercept. Only regression parameters are fitted; both methods predict one mean profile per perturbation. Resources: https://huggingface.co/ctheodoris/Geneformer and https://github.com/bowang-lab/scGPT.

##### State configurations

Figure 2D applies the released ST-HVG-Replogle checkpoint fewshot/hepg2/checkpoints/best.ckpt to all three genetic benchmarks without further fitting [15]. It uses Replogle/Nadig data with the released HepG2 split, 2,000-HVG expression inputs and one-hot perturbations. The released configuration specifies energy-distance loss, learning rate 10^*™*3^, batch size 64, seed 42 and up to 100,000 steps.

Figure 2E uses released ST-HVG-Tahoe zero-shot predictions for C32, HOP62, HepG2-C3A, Hs 766T and PANC-1, with training on the remaining Tahoe lines. Predictions are converted from log1p(CP1872) to log1p(CP10K); unmodeled genes retain matched control expression. Releases: https://huggingface.co/arcinstitute/ST-HVG-Replogle and https://huggingface.co/arcinstitute/ST-HVG-Tahoe.

##### X-Cell training

Open-X-Cell [14] has approximately 60 million parameters, 12 layers and hidden size 512, initialized from human scGPT gene embeddings and transformer weights [29]. Training uses six epochs on X-Atlas/Orion HCT116/HEK293T, followed by 20 epochs on Replogle/Nadig K562/RPE1/HepG2/Jurkat with 2,000 measured genes. The final checkpoint.pt predicts VCC/H1 and HepG2 responses using ESM-2, STRING, GenePT, DepMap and Cell Painting priors. HCT116 is excluded because all 207 test targets appeared in pretraining. Resources: https://github.com/zongtingwei/Open-X-cell and https://huggingface.co/zongting/Open-X-Cell.

##### PIE selection and inference

Figure 2D uses released PIE_replogle_wdataset checkpoints: rpe1/best_auprc.ckpt for VCC/H1, hepg2/best_auprc.ckpt for HepG2 and jurkat/best_auprc.ckpt for HCT116 [13]. Each folder names the held-out line; training and validation use the other three among HepG2, Jurkat, K562 and RPE1. Released checkpoints are selected by validation binary AUPRC. HepG2 uses its held-out fold.

For VCC/H1 and HCT116, all four folds are compared on 44 and 41 validation perturbations disjoint from testing. With seed 1, validation samples up to 512 cells per perturbation and 512 controls, scoring 17,502 and 18,530 uniquely mapped measured genes shared by all folds. HCT116 validation uses the shared QC dataset. Selection minimizes mean rank across PES, ΔPearson and ΔPert *r*, ranking higher scores first and breaking ties in the order HepG2, Jurkat, K562, RPE1. RPE1 is selected for VCC/H1 and Jurkat for HCT116, where Jurkat ties with K562. These configurations supply all four test metrics and remain fixed across full-gene and HVG panels.

PIE uses control expression, biological priors and released training-response evidence without local fitting. It predicts mean expression changes; MMD uses populations reconstructed from controls. Resources: https://huggingface.co/arcinstitute/PIE_replogle_wdataset and https://github.com/ArcInstitute/pie.

##### Stack inference

Figure 2F uses frozen Stack-Large-Aligned/bc_large_aligned.ckpt with ICL_FinetunedModel [16]. The model card reports ten pretraining epochs on human scBaseCount and eight alignment epochs on CellxGene and Parse PBMC. Context cells are pooled by cell type and cytokine from PBMC Donors 1, 3, 5, 6, 7, 9, 11 and 12; predictions transform controls from test Donors 2 and 8. Generation uses five masked-generation steps, prompt ratio 0.25, context ratio 0.4 (minimum 0.2), batch size 16 and seed 0. Inputs undergo expm1; nonnegative predicted counts are returned to log1p expression. Resources: https://huggingface.co/arcinstitute/Stack-Large-Aligned and https://github.com/ArcInstitute/stack.

##### PerturbMean baseline

Figure 2D assigns every test perturbation the cell-count-weighted mean expression of the corresponding benchmark’s genetic training conditions. Panel E averages the same drug-dose condition across Tahoe training lines; panel F averages the same cytokine response within each cell type across eight training donors. Missing matches use the global training mean. Distributional evaluation treats these means as point predictions.

###### HepG2 perturbation selection and visualization

For each of 258 held-out HepG2 perturbations, the mean response across 9,191 genes is centered on 512 matched controls: 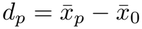. Greedy farthest-point sampling starts with the largest response norm, then maximizes the minimum Euclidean distance to selected responses:

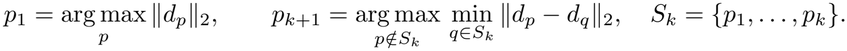

The ten selected targets are EIF2S1, UTP15, DHX16, RUVBL2, UBAP1, GMPPB, PAF1, GSPT1, SF3B5 and FAU. Observed, UPM and X-Cell panels use matched cell counts per target, capped at 256 and limited by availability, totaling 542 perturbed cells each. PCA (50 components) and UMAP (15 neighbors, minimum distance 0.1, seed 42) are fitted to these observed cells and 512 controls, then applied to predictions. Panels share axis limits and equal coordinate scaling. MMD averages full-population scores across the ten targets. Each target’s inter-perturbation distance is 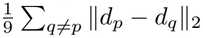, calculated identically for observed and predicted mean responses.

##### Extended HepG2 visualizations

Figures S2–S4 show nested sets of 10, 50 and all 258 held-out targets across 9,191 genes. The first set matches Figure 1D; continuing farthest-point sampling gives the 50-target set. All views reuse the Figure 1D PCA–UMAP reference, fitted to ten observed perturbations and 512 controls with 50 components, 15 neighbors, minimum distance 0.1 and seed 42. Shared observed, UPM and X-Cell coordinates match Figure 1D. UPM averages L6/L10 predictions; Gene-editing-only uses the same projection. Observed, UPM, Gene-editing-only and X-Cell each display 542, 2,568 and 17,589 perturbed cells in the 10-, 50- and 258-target views, respectively, plus 512 controls. Shared targets retain cells and colors, panels share axis limits, and MMD uses full-gene evaluation populations for each displayed target.

GEARS [27] adds predicted mean changes to sampled controls and clips negative values to zero, using the validation-selected epoch 2 checkpoint from 20 epochs of training. Its GO mapping covers 9, 49 and 255 targets, with 468, 2,494 and 17,273 displayed cells. FAU is absent from the 10- and 50-target sets; C19orf53, C7orf26 and FAU are absent from the 258-target set. Covered targets use matched cell counts; MMD averages only covered targets. Projected cells can overlap.

##### UPM ensemble

L6 and L10 are six-layer and ten-layer UPM models trained on the final collection. Figure S5 compares L10 with their equal-weight ensemble on the Figure 2D subsets: 29 VCC/H1, 258 HepG2 and 207 HCT116 targets across 17,540, 9,191 and 18,907 genes. Ensembling lowers MMD and raises both response correlations on all three datasets; PES rises on HepG2/HCT116 and falls on VCC.

##### Evaluation gene-panel sensitivity

Figures S6–S11 vary evaluation genes while retaining checkpoints and test perturbations. GEARS uses the same validation-selected VCC/H1 epoch 1 and HepG2 epoch 2 checkpoints from 20-epoch runs as Figure 2D. All four metrics are recomputed on full-gene, HVG200 and HVG2000 panels. HVGs are ranked separately per dataset by single-cell variance in training-only log1p(CP10K) expression, including controls. The top 200 are nested within the top 2,000, with ties resolved by original column order.

PIE retains the configurations in Appendix B.4: RPE1 for VCC/H1, HepG2 for HepG2 and Jurkat for HCT116. Expanded comparisons add the UPM ensemble and alternative PIE configurations. Auxiliary PIE MMD scores are shown but excluded from best-score highlighting. Missing outputs receive zero expression change; GEARS and X-Cell are omitted from HCT116. Model resources follow the main comparison [13, 14, 15, 27, 28, 29].

##### Mean-response heatmaps and UMAP

Figures S12–S17 retain the Figure 2D targets, measured genes and standalone L10 UPM. Heatmap rows follow average-linkage clustering of observed responses; columns follow decreasing mean absolute observed response. A shared symmetric color scale is capped at the 99th percentile of absolute observed changes. Mean-response UMAP jointly embeds observed and predicted control-centered profiles using up to 50 principal components, 15 neighbors, minimum distance 0.1, Euclidean distance and seed 42. Exact duplicates are collapsed for fitting and receive identical coordinates. Each point represents one perturbation mean.

##### Extended perturbation sets

Figures S18–S23 cover 93 H1, 476 HepG2 and 228 HCT116 perturbations on reference panels of 17,540, 9,191 and 18,827 genes. They include standalone UPM and its two-model ensemble, omit PIE, and retain earlier Geneformer outputs on 1,387 genes. GEARS uses the same validation-selected 20-epoch runs as Figure 2D, covering 29 H1 perturbations on 17,520 genes and 255 HepG2 perturbations on 9,189 genes; HCT116 omits GEARS. Gray heatmap regions indicate unavailable predictions. Heatmap rows follow alphabetical perturbation order, and columns are ordered by observed-change variance. Colors represent control-centered mean log1p(CP10K) expression on a shared symmetric scale within each dataset. Extended mean-response UMAP uses 1,387 shared genes, with PCA (30 components) and UMAP (15 neighbors, minimum distance 0.3, seed 42) fitted to observed responses only; predictions and controls are projected into this reference.

##### B.4.1 Combination evaluation

###### Combination evaluation set

Gene-gene evaluation uses 131 Norman pairs and 16,185 measured genes, with both singleton perturbations represented in training [20]. Gene-drug evaluation uses 12,083 sci-Plex-GxE conditions and 18,904 genes across A172, T98G and U87MG [30]. Supplementary Figure S24 evaluates seven held-out SciPlexCombo/A549 pairs with both singleton responses available [24]; Figure 2B uses the broader 25-combination set.

###### Combination baselines

All combination responses are expressed in log1p(CP10K) space. For constituent perturbations

*A* and *B*, the observed-singleton Additive baseline is

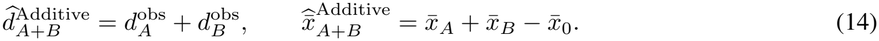

Here 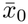 is the matched control mean. Additive requires measured singleton responses at prediction time. PerturbMean assigns the mean single-perturbation expression within the same cellular context. MMD treats both as point distributions.

Gene-only + Drug-only sums independently predicted changes for genetic target *g* and drug-dose input (*m, a*) using the same unperturbed cell *x*_0*i*_:

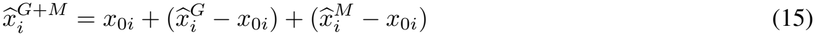

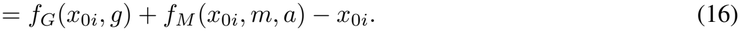

Gene-editing-only and Molecule-only receive genetic and drug-dose inputs, respectively; UPM receives both. Their predicted changes are summed without clipping or renormalization. All native-input comparisons use matched deterministic control samples, capped at 32 cells per condition for gene-drug MMD.

The Additive-assisted Ensemble is

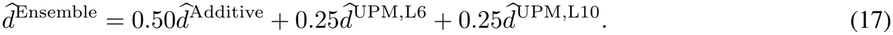

Both UPM configurations use the final training collection.

###### Combination performance metrics

Following Appendix B.3, ΔPearson correlates responses across genes within each combination, and ΔPert *r* correlates responses across combinations for each gene. Norman has 16,173 finite gene-wise scores. MMD compares cell populations before dimensionality reduction.

###### Non-additivity and win fractions

Combination difficulty is the retrospective cosine distance between observed combination and observed-singleton Additive responses. For nonzero vectors *a* and *b*,

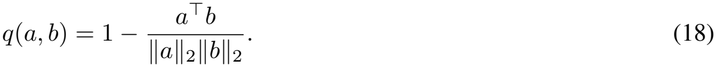

Vectors span genes for condition-wise difficulty and conditions for gene-wise difficulty. MMD uses condition-wise difficulty. Comparisons retain only finite, eligible scores. In bin B, the win percentage of predictor *A* over *B* is

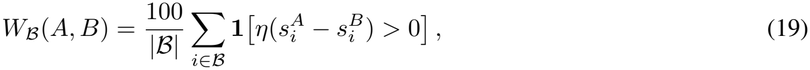

where *η* = 1 for correlations and *η* = ™1 for MMD; ties do not count as wins. Histograms report eligible genes or conditions per bin, and overall wins use all eligible observations. The merged Norman condition-wise tail at 0.35–0.60 contains six pairs. Uniform zero or 100% win bars may be omitted, while model score curves are retained.

##### Drug-drug evaluation

Figure S24 applies the same cosine distance to seven SciPlexCombo/A549 combinations, with controls pooled across A549 wells. Of 15,735 output genes, 15,728 have finite correlations per predictor. Condition-wise and gene-wise bins have widths 0.05 and 0.1. Lines join descriptive bin means; lower bars show pairwise wins, with counts reported for the seven evaluated combinations. In this drug-drug comparison, Molecule-only and UPM receive chemical inputs.

##### Genetic-target specificity diagnostic

Each standalone UPM gene-drug prediction is replaced by the mean within its cell-line/drug/dose group, removing genetic-target variation while preserving chemical and cellular-context responses. Global gene-wise correlation was maintained after replacement.

##### B.4.2 HSC application evaluation

###### HSC inputs and experimental references

Both HSC settings use human MASH expression states and published response directions. Following Kim et al. [17], GAS7, SERPINE1, SPON1 or LTBP2 knockdown is combined with TGF*β*1, with ACTA2, COL1A1, SERPINE1 and LOXL2 readouts. Following Liu et al. [18], ETS1, ETS2, GATA4 or GATA6 knockdown uses COL1A1, LOXL2, IL6 and CXCL5 readouts. Each reference setting contains 16 target-readout pairs.

###### HSC prediction settings

UPM receives knockdown alone or with TGF*β*1. Gene-editing-only uses the epoch 2,000 checkpoint from Figure 2B and receives knockdown only in both settings. All predictions use 1,152 human HSC cells, 128 from each of nine donors. For normalized log-expression *x*_*i*_, genetic predictor *f*_*G*_ and target *t*,

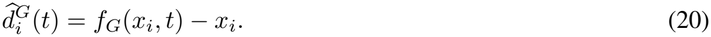

Predictors retain their native output normalization in log1p(CP10K) space. Figure 4B compares Gene-editing-only with joint-input UPM; Figure 4C compares Gene-editing-only, X-Cell and UPM on single knockdowns. HSC X-Cell uses the six-epoch Orion-pretrained Open-X-Cell Mini-model [14]. Resources: https://github.com/zongtingwei/Open-X-cell and https://huggingface.co/zongting/Open-X-Cell.

###### Random-knockdown normalization

For each target *t* and readout *g*, predicted changes are averaged over 128 cells per donor, then across nine donors, yielding 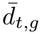. The same procedure is applied to 64 fixed random-knockdown targets *ℛ*. Normalized scores are

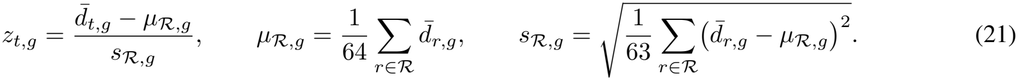

Positive and nonpositive scores classify upregulation and downregulation, respectively. In the cytokine setting, UPM uses matched combined interventions for both targets and random-knockdown references. Gene-editing-only uses knockdown-only predictions throughout. Scores measure target-relative effects within each predictor.

###### Response-direction metrics

For response-direction class *k* ∈ {up, down}, let *TP*_*k*_, *FP*_*k*_ and *FN*_*k*_ denote true positives, false positives and false negatives. The class and macro scores are

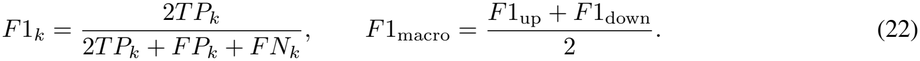

For the *T* = 32 target-readout pairs with reference directions *y* and predicted directions *y*,

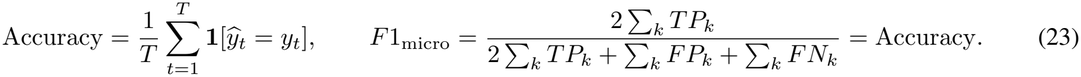

Macro-F1 averages the up/down class scores; micro-F1 equals accuracy for this single-label binary task. The human combined-intervention group has 16 down reference directions, and the mouse-reference single-KD group has 16 up directions. Group accuracies are reported separately; the pooled set is class-balanced. Gene-editing-only covers both groups, and X-Cell only single knockdowns.

**Table S2:** Directional prediction performance across 32 HSC target-readout pairs.

| Method | Macro-F1 | Micro-F1 | Correct / 32 |
|---|---|---|---|
| All-down | 0.3333 | 0.5000 | 16 |
| All-up | 0.3333 | 0.5000 | 16 |
| Gene-editing-only | 0.4640 | 0.4688 | 15 |
| UPM | 0.6532 | 0.6562 | 21 |

Table S2 reports 6/16 and 9/16 correct directions for Gene-editing-only in the human combined-intervention and mouse-reference single-KD groups, versus 9/16 and 12/16 for UPM. X-Cell recovers 7/16 in single-KD evaluation. Pooled UPM accuracy is 21*/*32 = 65.625%, versus 15*/*32 = 46.875% for Gene-editing-only: a (21*/*15 ™ 1) × 100% = 40.0% relative gain or 18.75 percentage points.

### C Additional results

**Figure S2:**
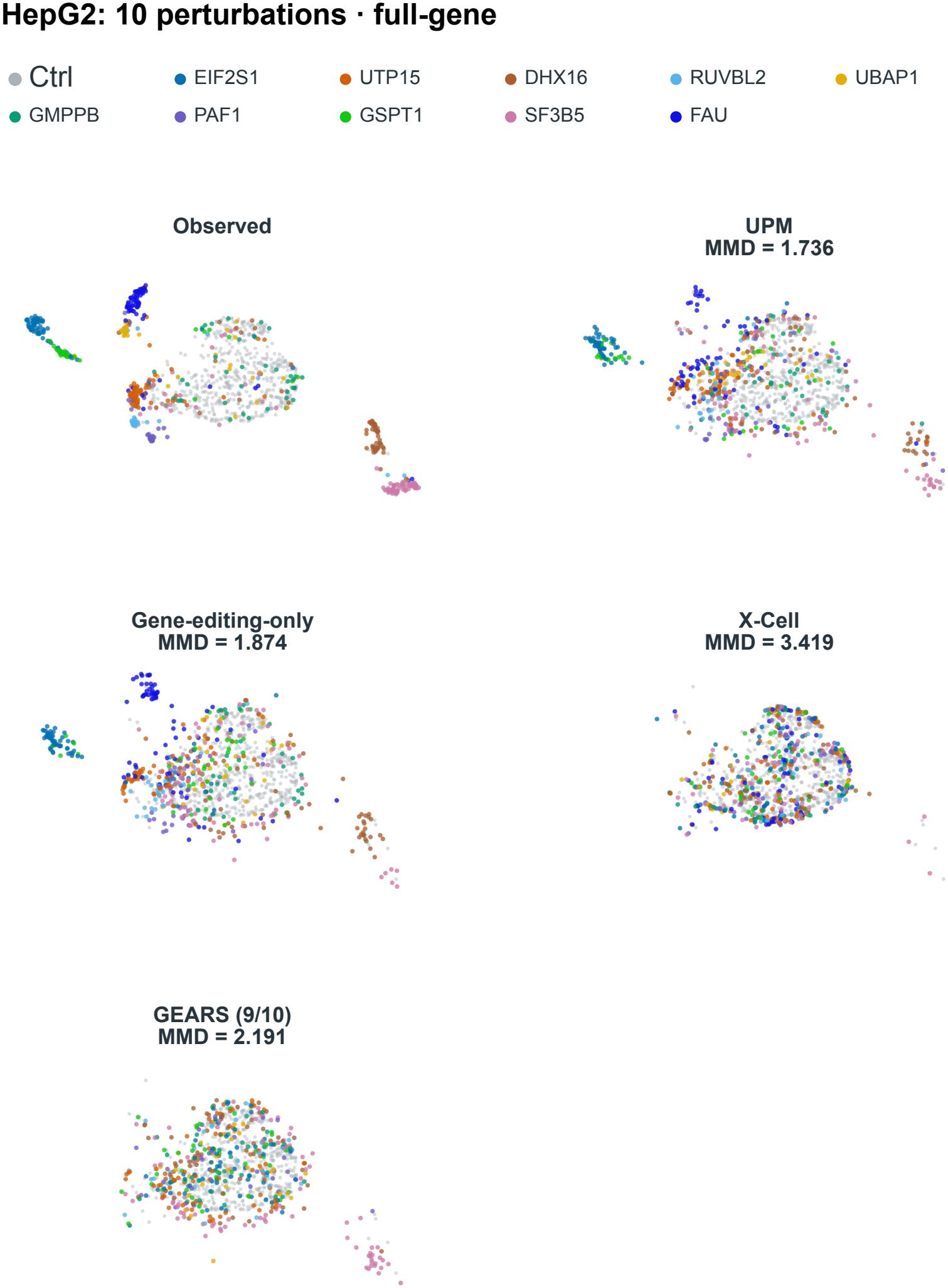
Single-cell UMAP of 10 held-out HepG2 perturbations across all measured genes. Observed and predicted cells are projected using the Figure 1D reference derived from 9,191 genes. Colors identify perturbations, and gray denotes control cells. Figures S2–S4 share the projection, axis limits and colors for common perturbations. UPM denotes the ensemble of L6 and L10 predictions. GEARS covers nine of the ten targets, excluding FAU; MMD is averaged over the targets displayed for each method. Cell sampling, projection parameters and checkpoint selection are described in Appendix B.4.

**Figure S3:**
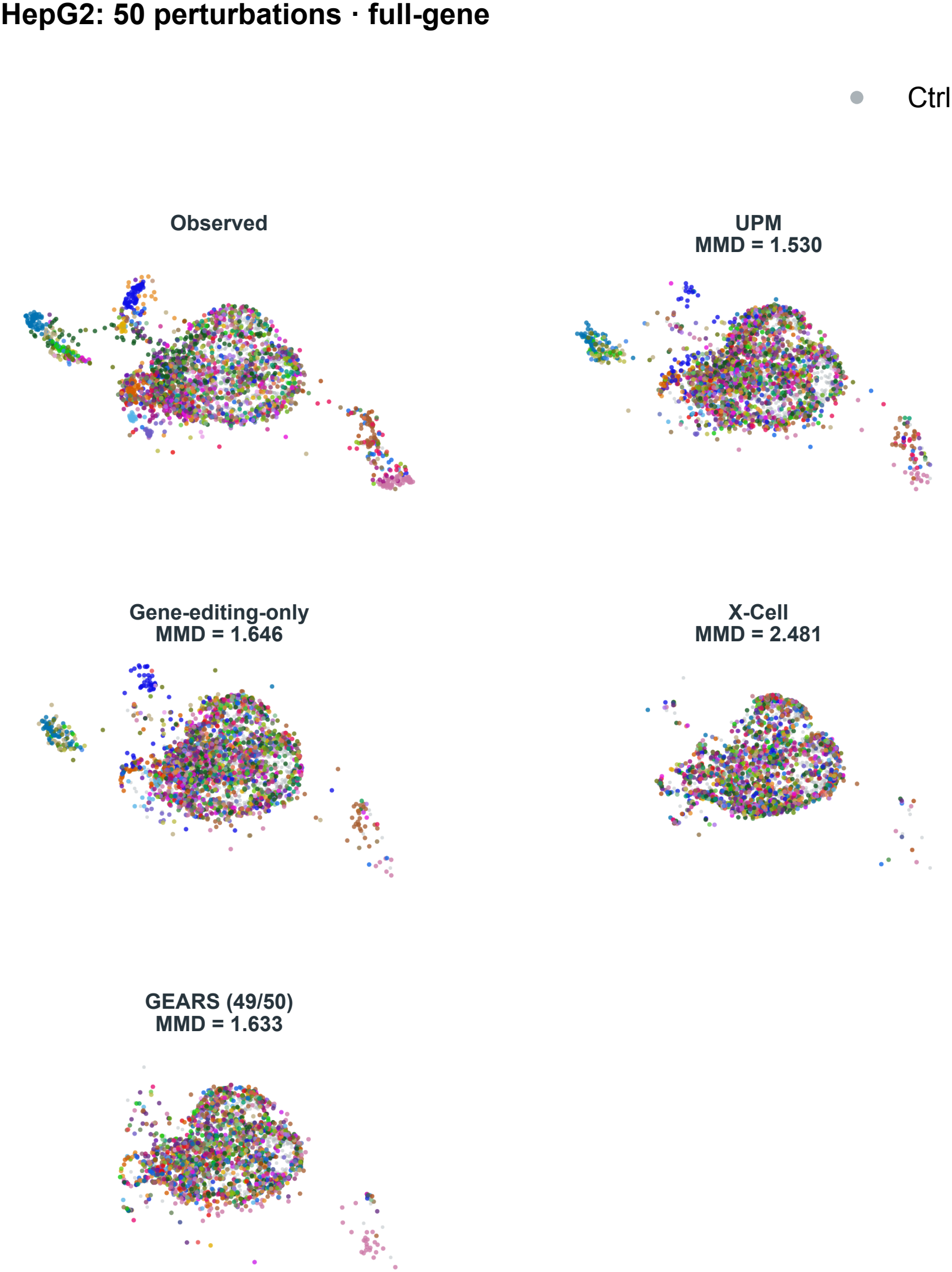
Single-cell UMAP of 50 held-out HepG2 perturbations across all measured genes. This subset includes the ten perturbations in Figure S2 and extends the comparison to 50 targets using the same 9,191-gene projection and display conventions. GEARS covers 49 targets, excluding FAU; MMD is averaged over the targets displayed for each method. Perturbation selection and visualization methods are described in Appendix B.4.

**Figure S4:**
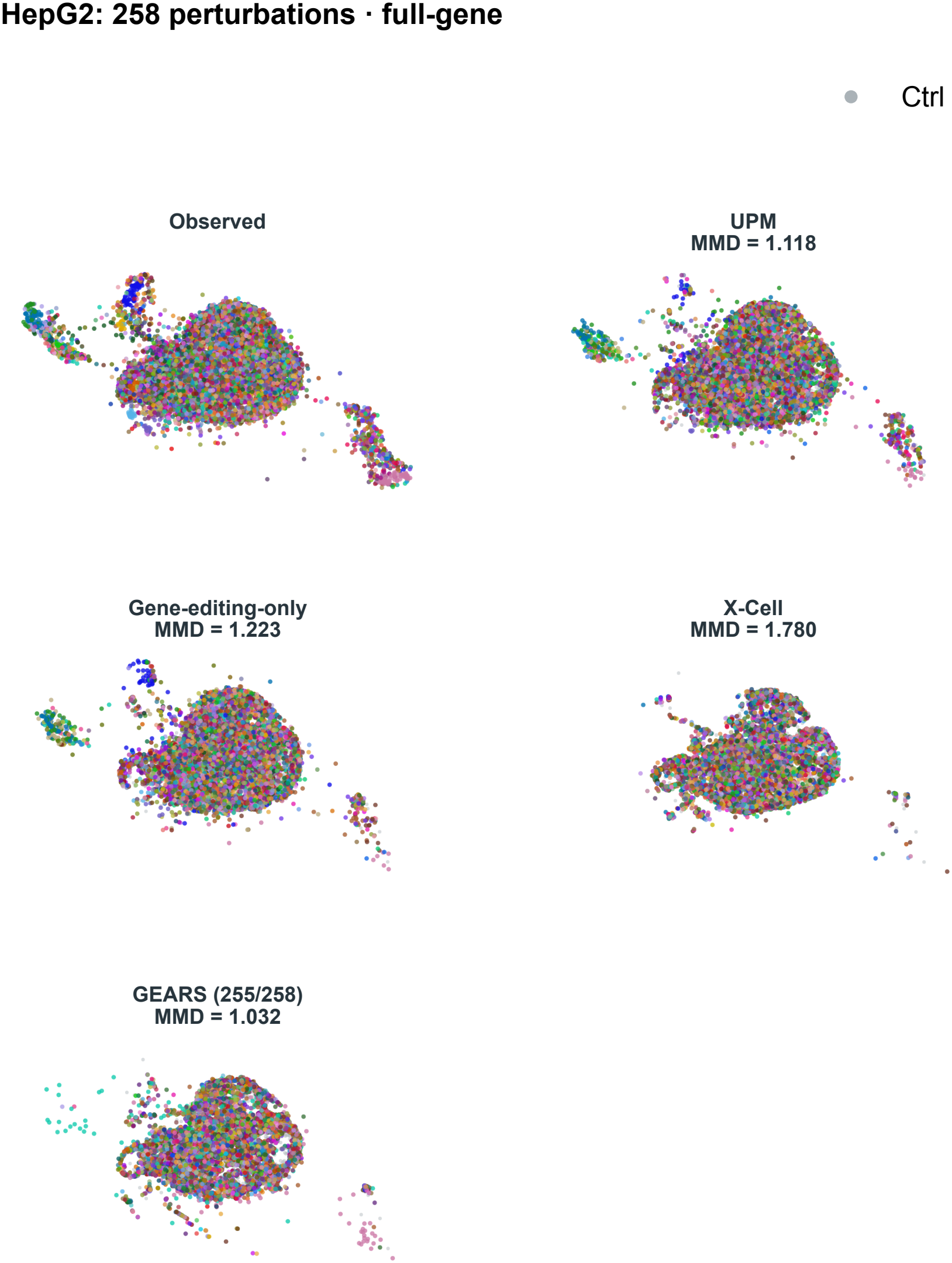
Single-cell UMAP of all 258 held-out HepG2 perturbations across all measured genes. This comparison includes the nested subsets in Figures S2 and S3, using the same 9,191-gene projection and display conventions. GEARS covers 255 targets, excluding C19orf53, C7orf26 and FAU; MMD is averaged over the targets displayed for each method. Visualization methods and model coverage are described in Appendix B.4.

**Figure S5:**
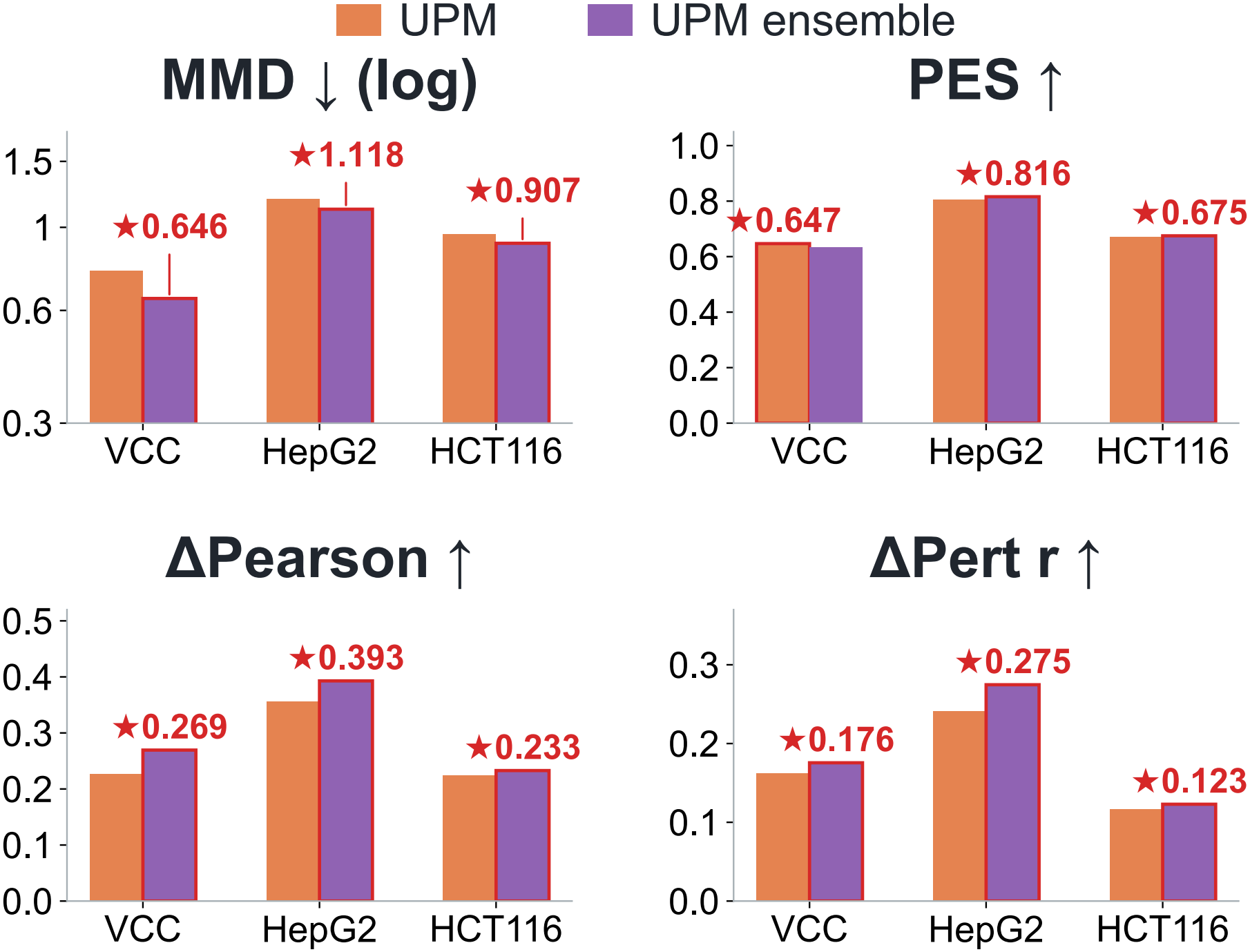
Comparison of standalone and ensemble UPM on genetic perturbation benchmarks. Standalone UPM uses L10, and the UPM ensemble averages L6 and L10 predictions. Evaluation uses the same perturbations and gene sets as Figure 2D. Bars show MMD, PES, ΔPearson and ΔPert *r*, with a logarithmic axis for MMD. Red outlines and stars identify the better-performing configuration for each metric.

**Figure S6:**
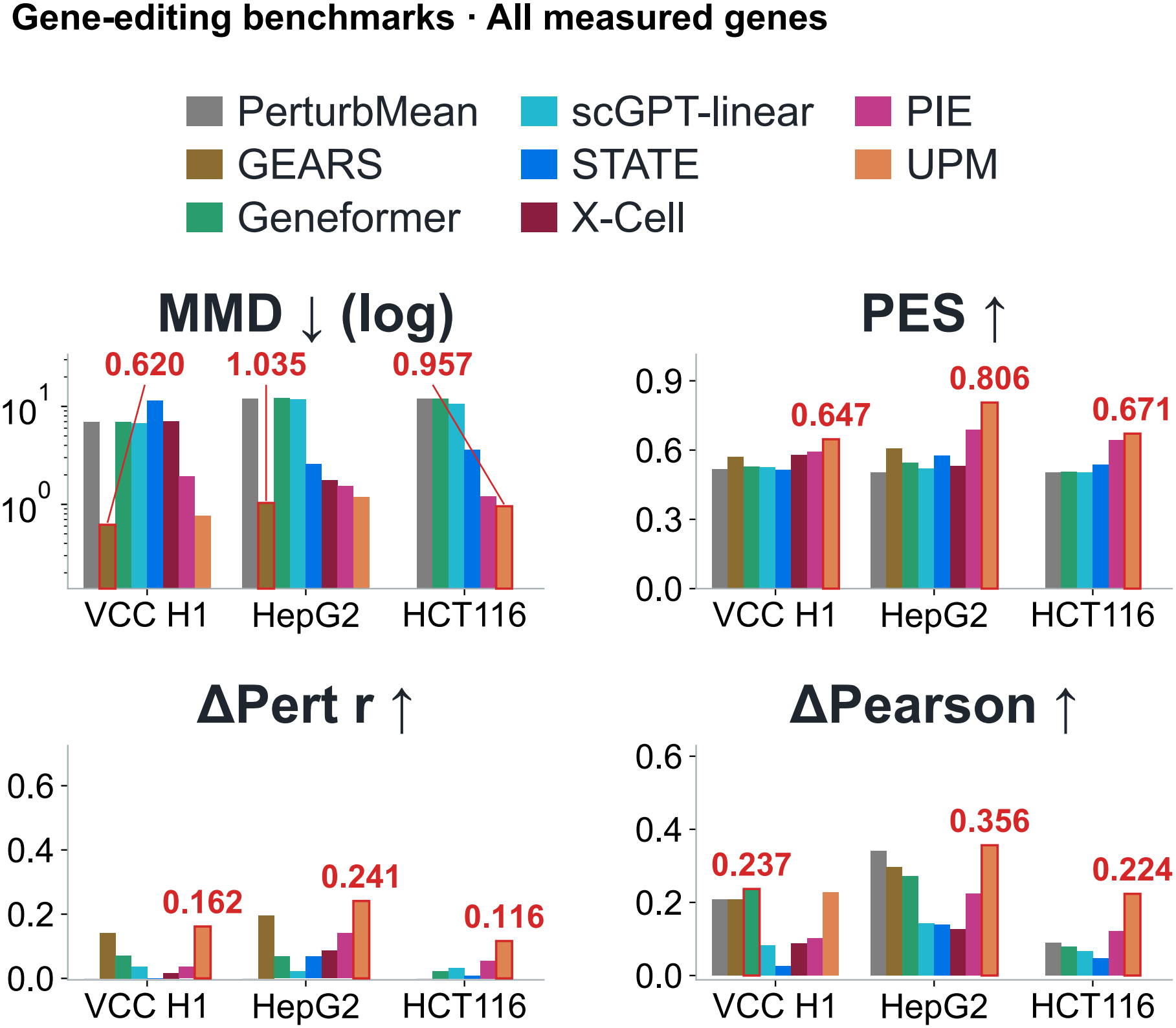
Model comparison on all measured genes. Evaluation uses the same test perturbations and measured gene sets as Figure 2D, with standalone L10 UPM. Figures S6–S8 compare evaluation gene panels while keeping model checkpoints and test perturbations fixed. Bars show MMD, PES, ΔPearson and ΔPert *r*, with a logarithmic axis for MMD. Red labels and outlines mark the best scores; PIE configurations are excluded only from MMD best-score selection. GEARS and X-Cell are shown only for VCC/H1 and HepG2. Model configurations and gene-panel selection are described in Appendix B.4.

**Figure S7:**
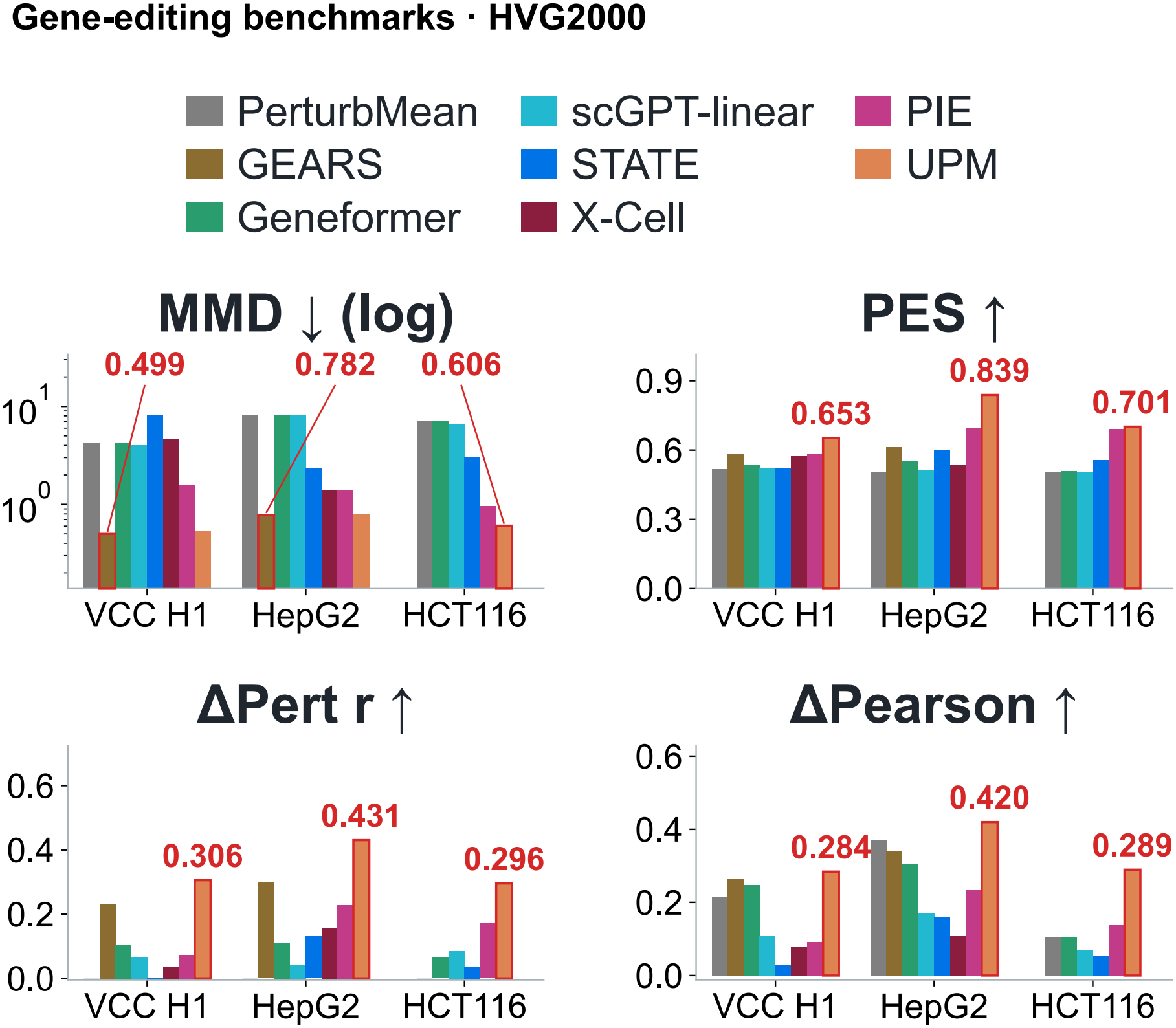
Model comparison on 2,000 highly variable genes. Models are evaluated on the same 2,000 HVGs selected from each dataset’s training data. Test perturbations, model checkpoints and display conventions follow Figure S6; only the evaluation gene panel changes.

**Figure S8:**
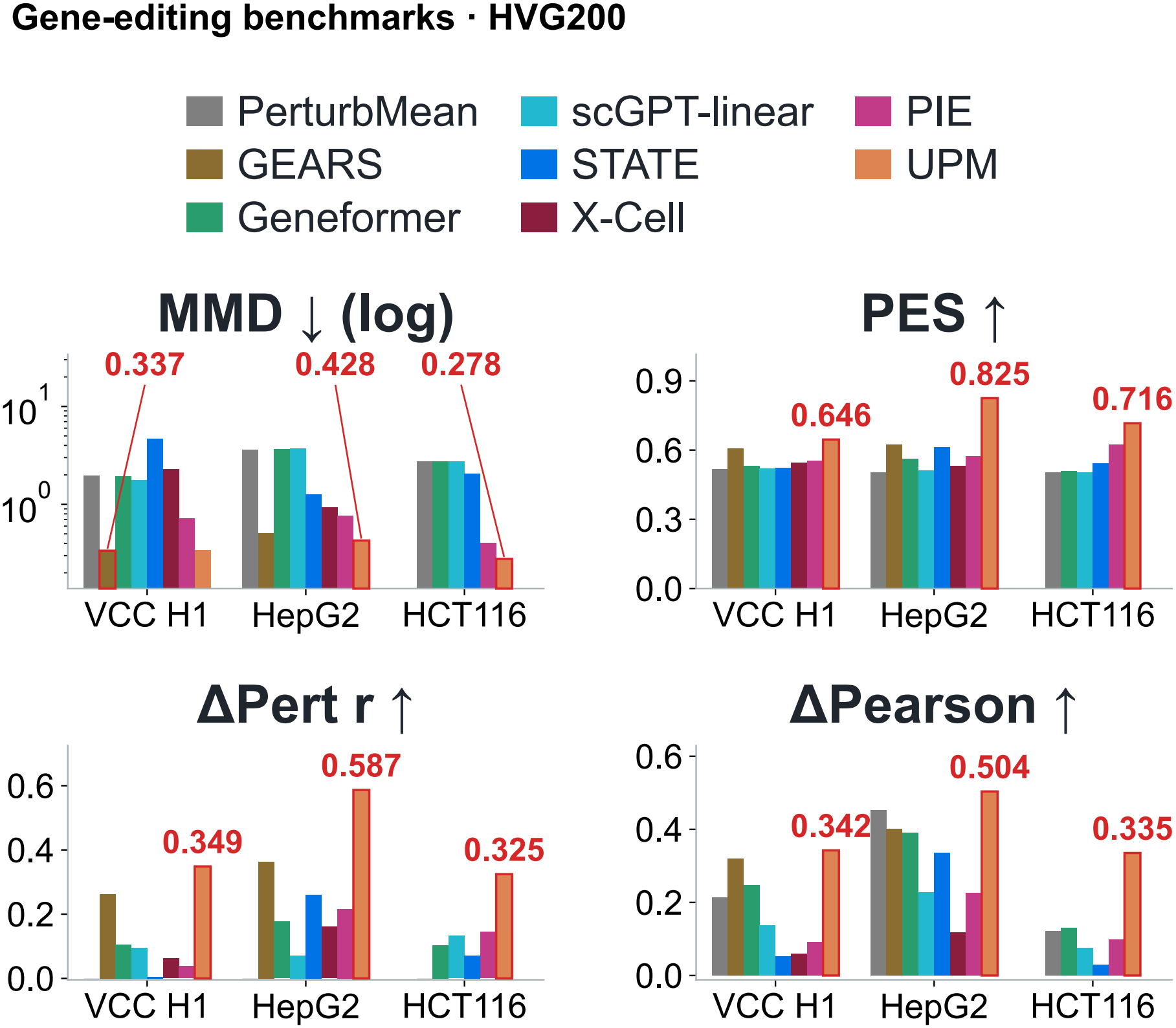
Model comparison on 200 highly variable genes. Models are evaluated on the top 200 training-derived HVGs within the 2,000-gene panels in Figure S7. Test perturbations, model checkpoints and display conventions follow Figure S6.

**Figure S9:**
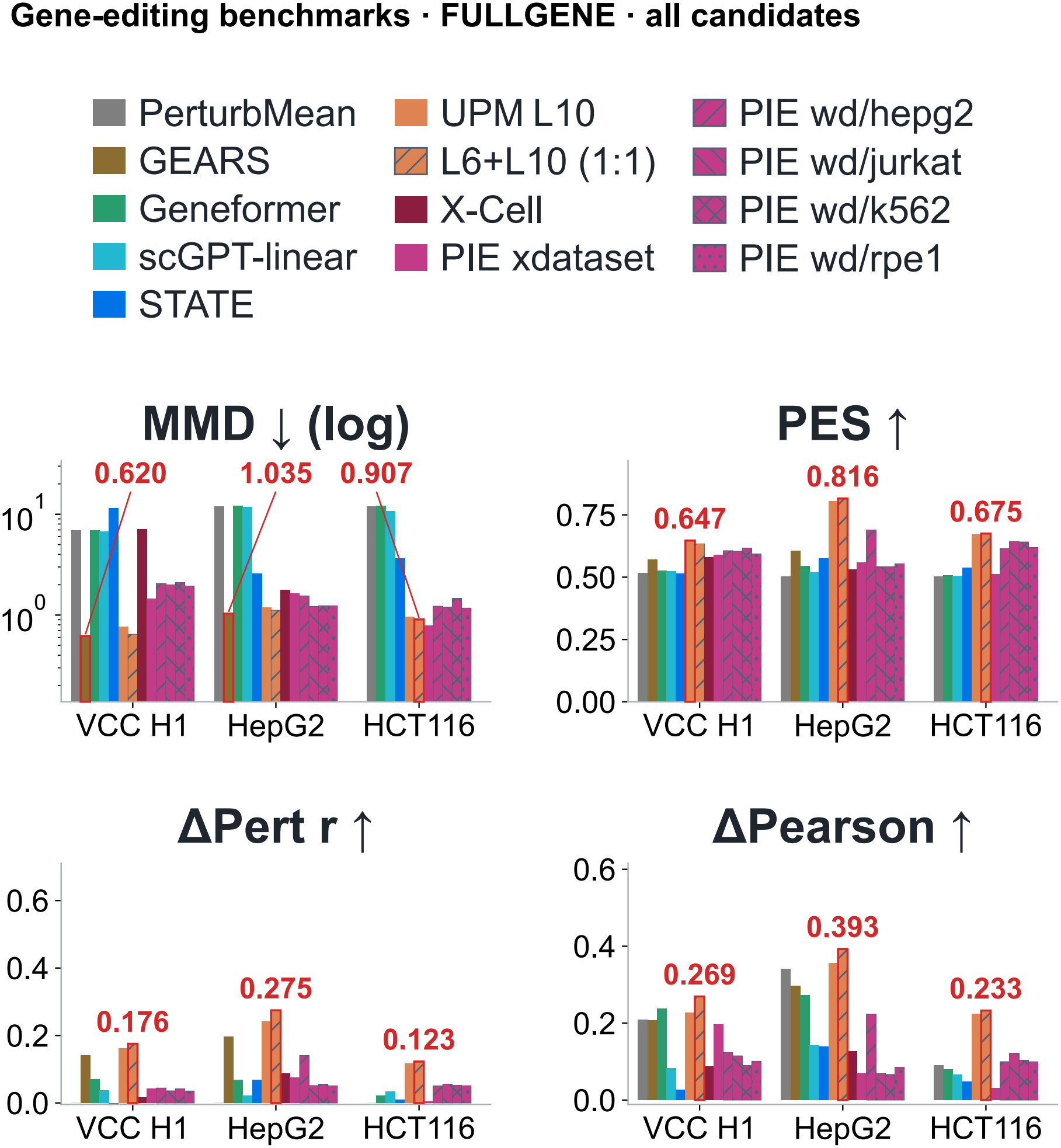
Expanded model comparison on all measured genes. This comparison extends Figure S6 by adding the UPM ensemble and four additional PIE configurations. The UPM ensemble averages L6 and L10 predictions with equal weights. Evaluation gene sets, test perturbations and display conventions follow Figure S6. PIE configurations are excluded only from MMD best-score selection; their definitions and selection are described in Appendix B.4.

**Figure S10:**
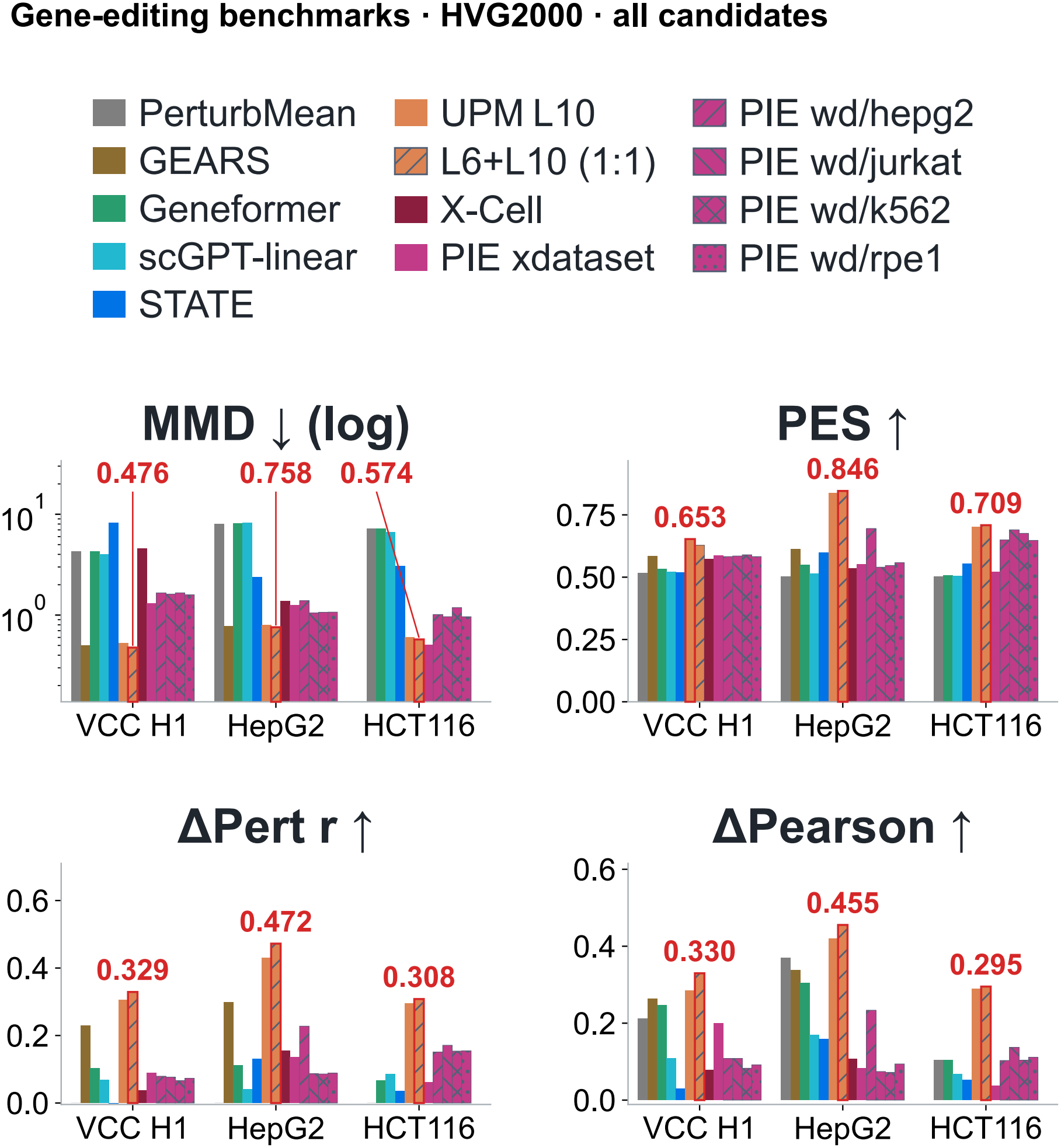
Expanded model comparison on 2,000 highly variable genes. The configurations in Figure S9 are evaluated on the same training-derived HVG2000 panels as Figure S7. Model checkpoints, test perturbations and display conventions remain unchanged.

**Figure S11:**
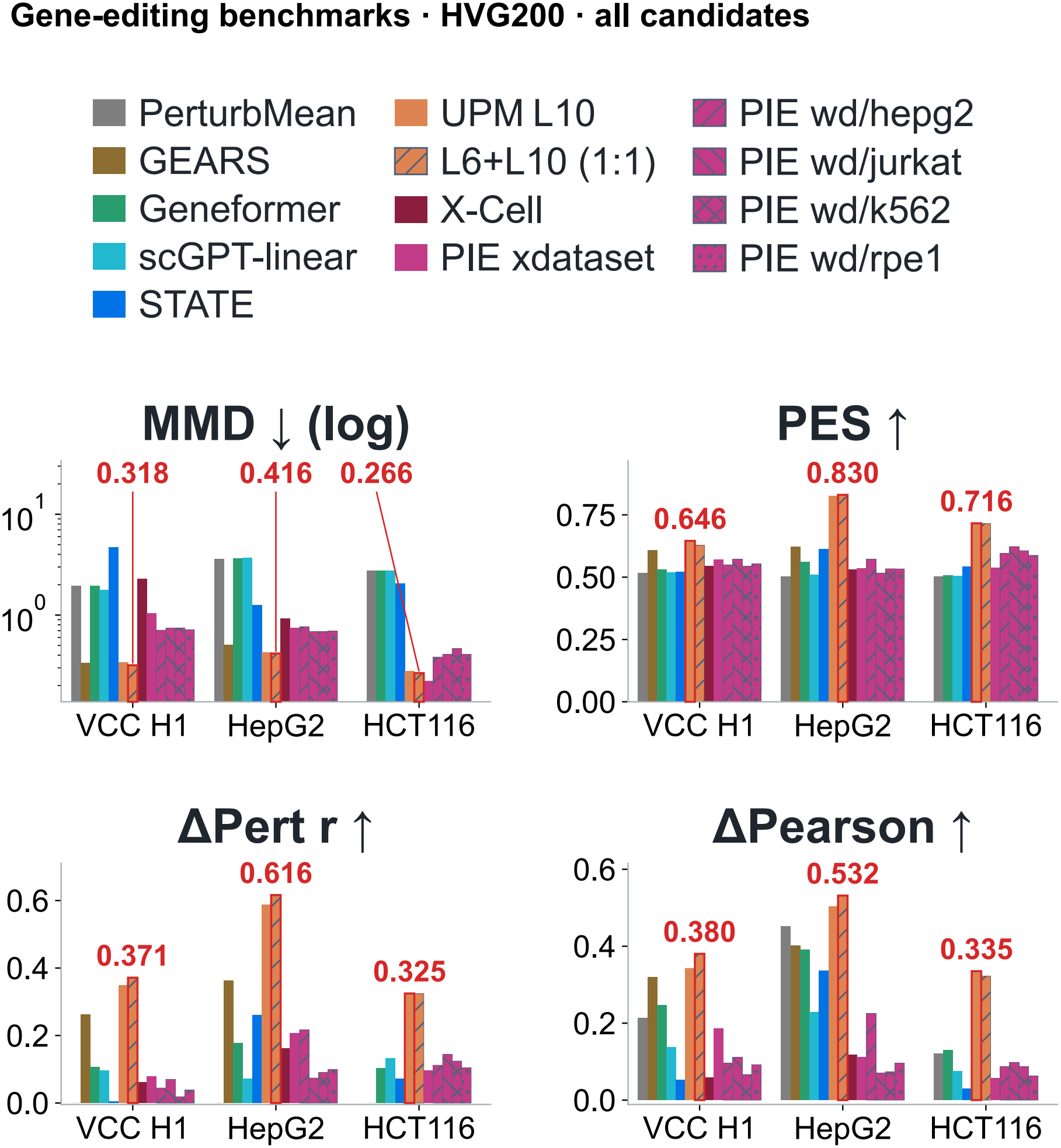
Expanded model comparison on 200 highly variable genes. The configurations in Figure S9 are evaluated on the same training-derived HVG200 panels as Figure S8. Model checkpoints, test perturbations and display conventions remain unchanged.

**Figure S12:**
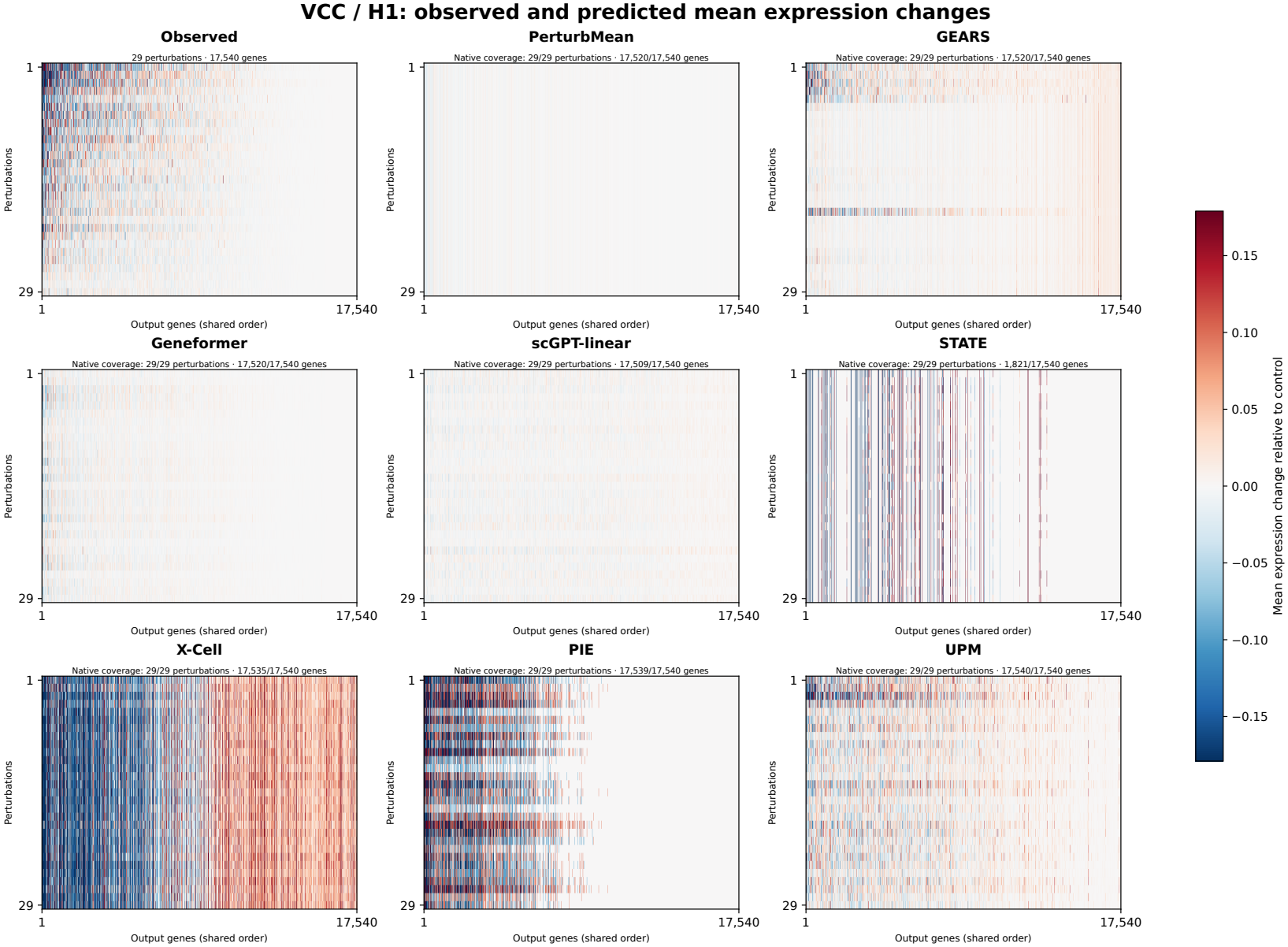
Observed and predicted mean responses on VCC/H1. Heatmaps compare observed responses with eight predictors across 29 perturbations and 17,540 genes, using the Figure 2D model configurations with standalone L10 UPM. Rows represent perturbations and columns represent genes, ordered identically across panels according to observed responses. Colors indicate mean expression changes relative to controls on a shared symmetric scale. Panel headers report perturbation and gene coverage. Ordering, model configurations and handling of missing outputs are described in Appendix B.4.

**Figure S13:**
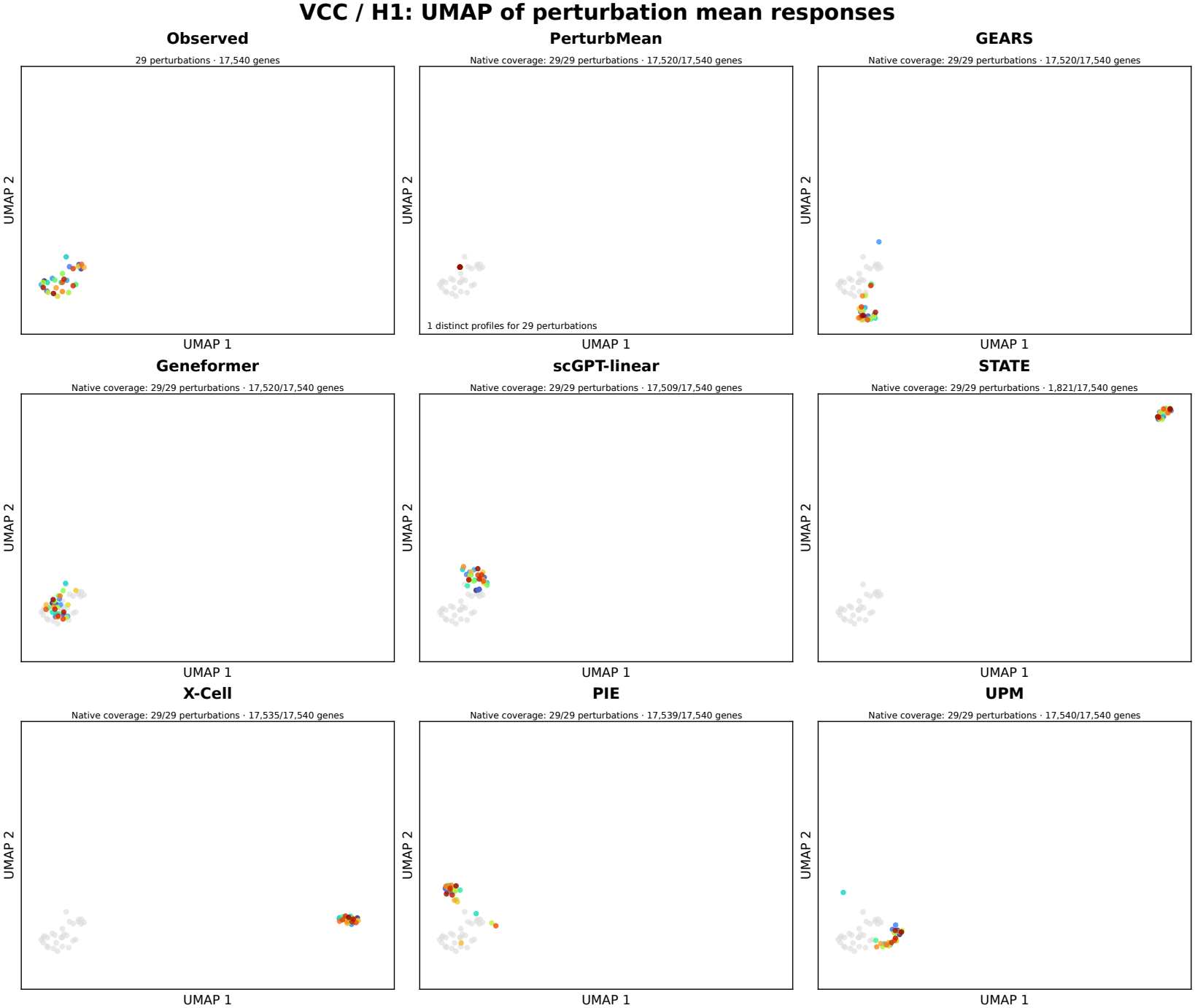
UMAP of perturbation mean responses on VCC/H1. Observed and predicted mean responses to 29 perturbations across 17,540 genes are jointly embedded. Each colored point represents a predicted perturbation response, and gray points show observed references. Panels share axis limits; PerturbMean predictions overlap because the same response is assigned to every perturbation. Model configurations match Figure S12, and embedding methods are described in Appendix B.4.

**Figure S14:**
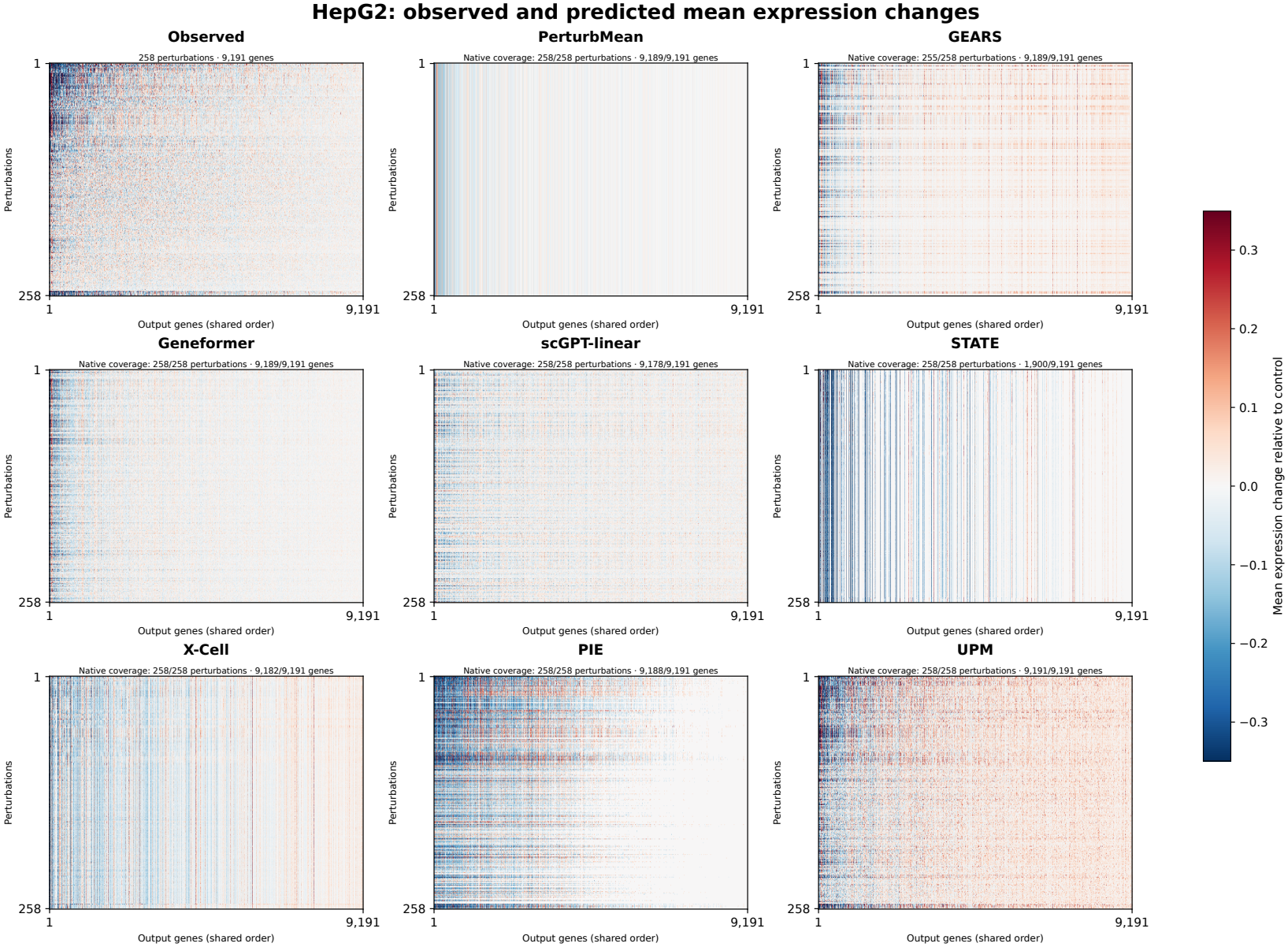
Observed and predicted mean responses on HepG2. Heatmaps compare observed responses with eight predictors across 258 perturbations and 9,191 genes, using the Figure 2D model configurations with standalone L10 UPM. Row and column ordering and color scaling follow the conventions in Figure S12, using HepG2 observed responses. Panel headers report perturbation and gene coverage.

**Figure S15:**
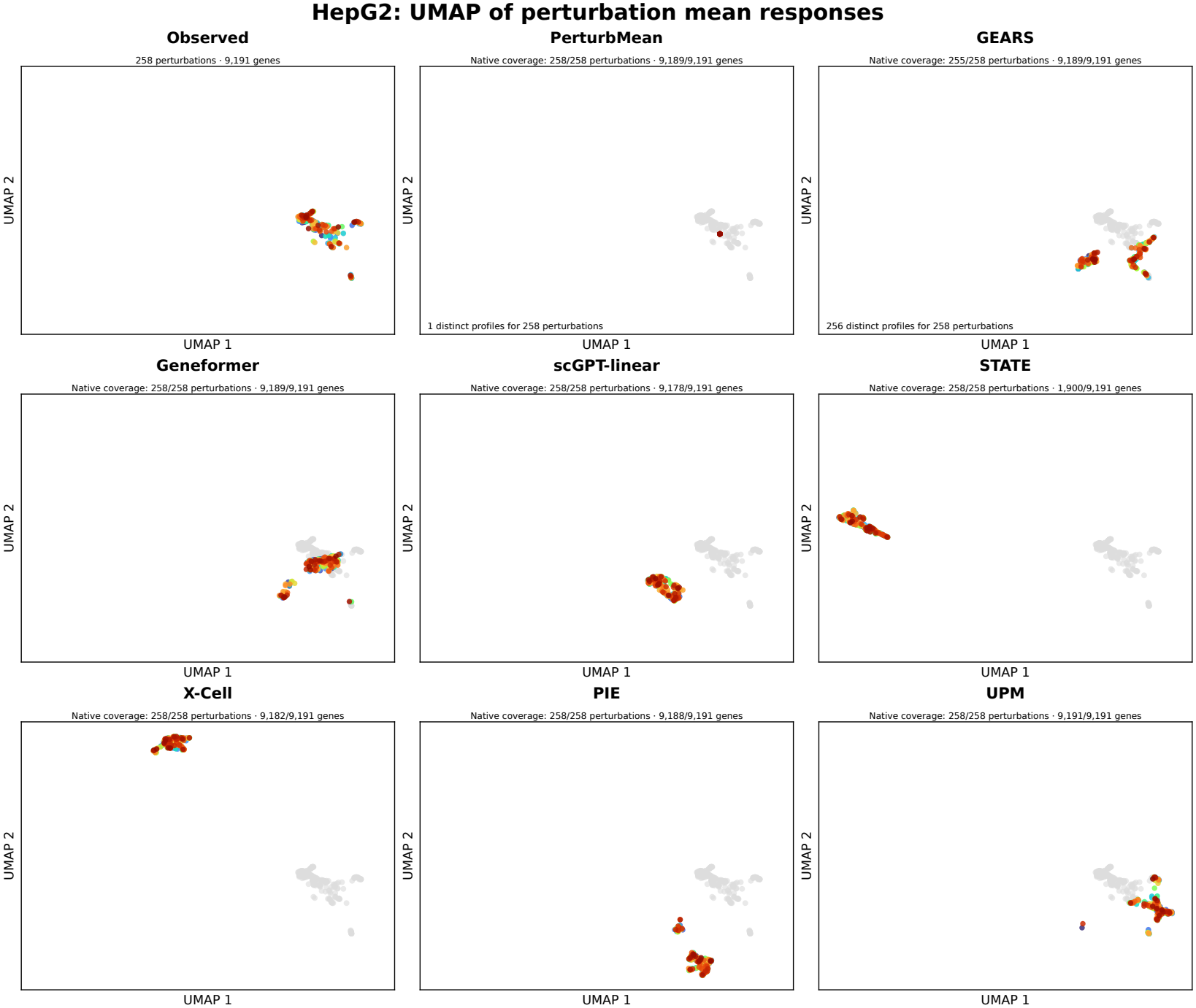
UMAP of perturbation mean responses on HepG2. Observed and predicted mean responses to 258 perturbations across 9,191 genes are jointly embedded, using the models in Figure S14. Colors identify predicted perturbation responses, and gray points show observed references. Panels share axis limits within each dataset. PerturbMean predictions overlap as described in Figure S13.

**Figure S16:**
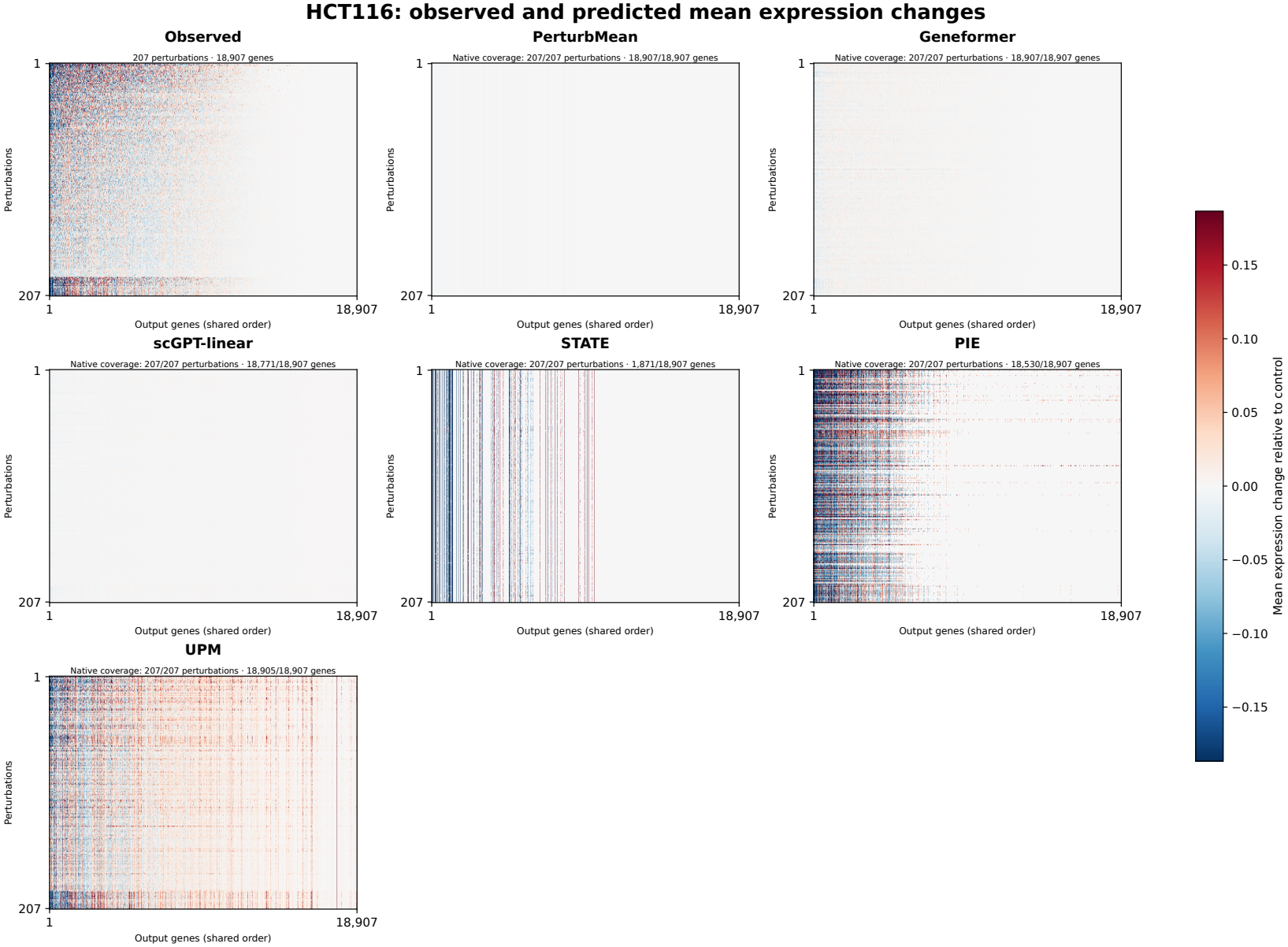
Observed and predicted mean responses on HCT116. Heatmaps compare observed responses with six predictors across 207 perturbations and 18,907 genes, using the Figure 2D model configurations with standalone L10 UPM. Row and column ordering and color scaling follow the conventions in Figure S12, using HCT116 observed responses. Panel headers report perturbation and gene coverage. GEARS and X-Cell are not included.

**Figure S17:**
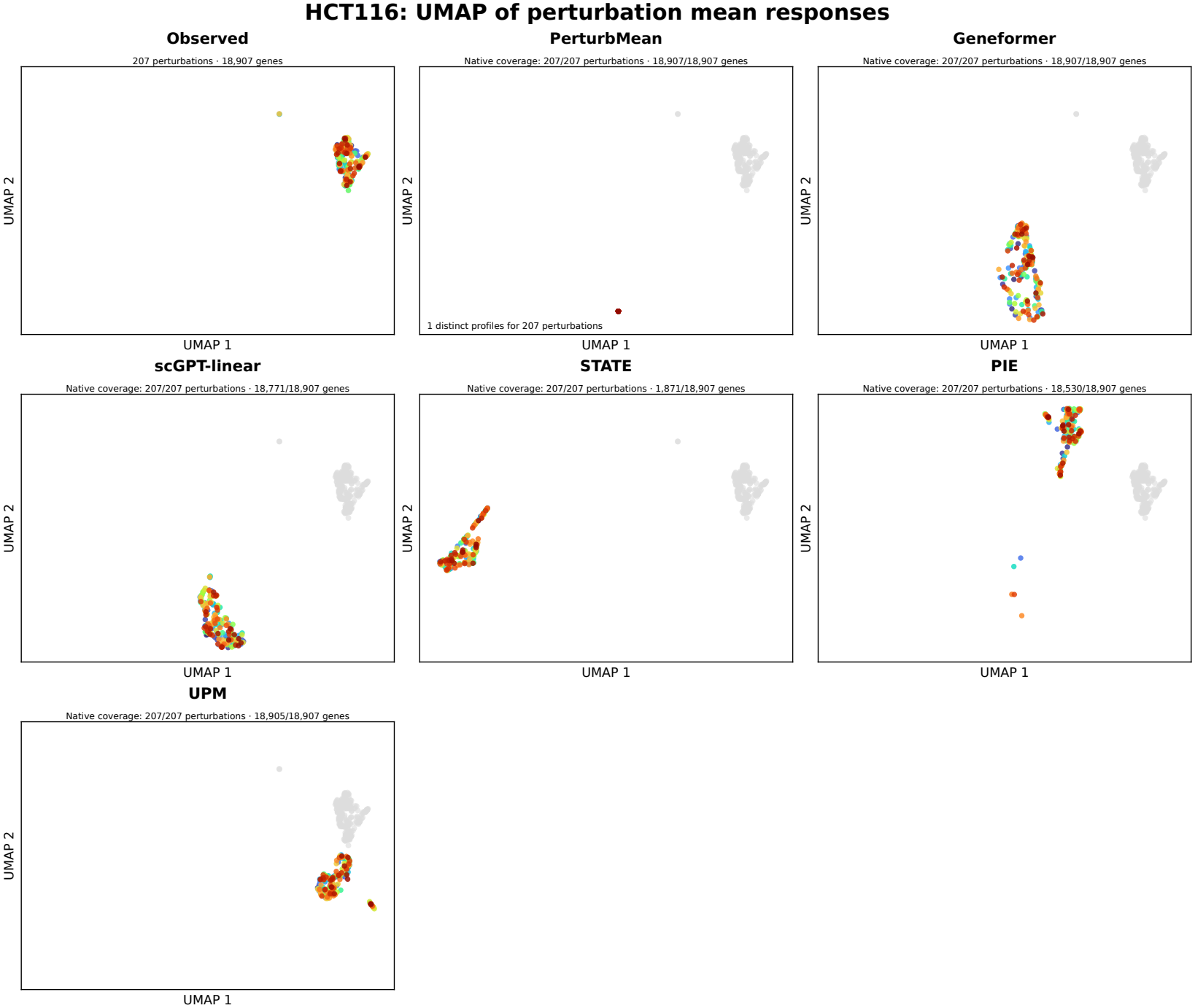
UMAP of perturbation mean responses on HCT116. Observed and predicted mean responses to 207 perturbations across 18,907 genes are jointly embedded, using the six models in Figure S16. Colors identify predicted perturbation responses, and gray points show observed references. Panels share axis limits within each dataset. PerturbMean predictions overlap as described in Figure S13.

**Figure S18:**
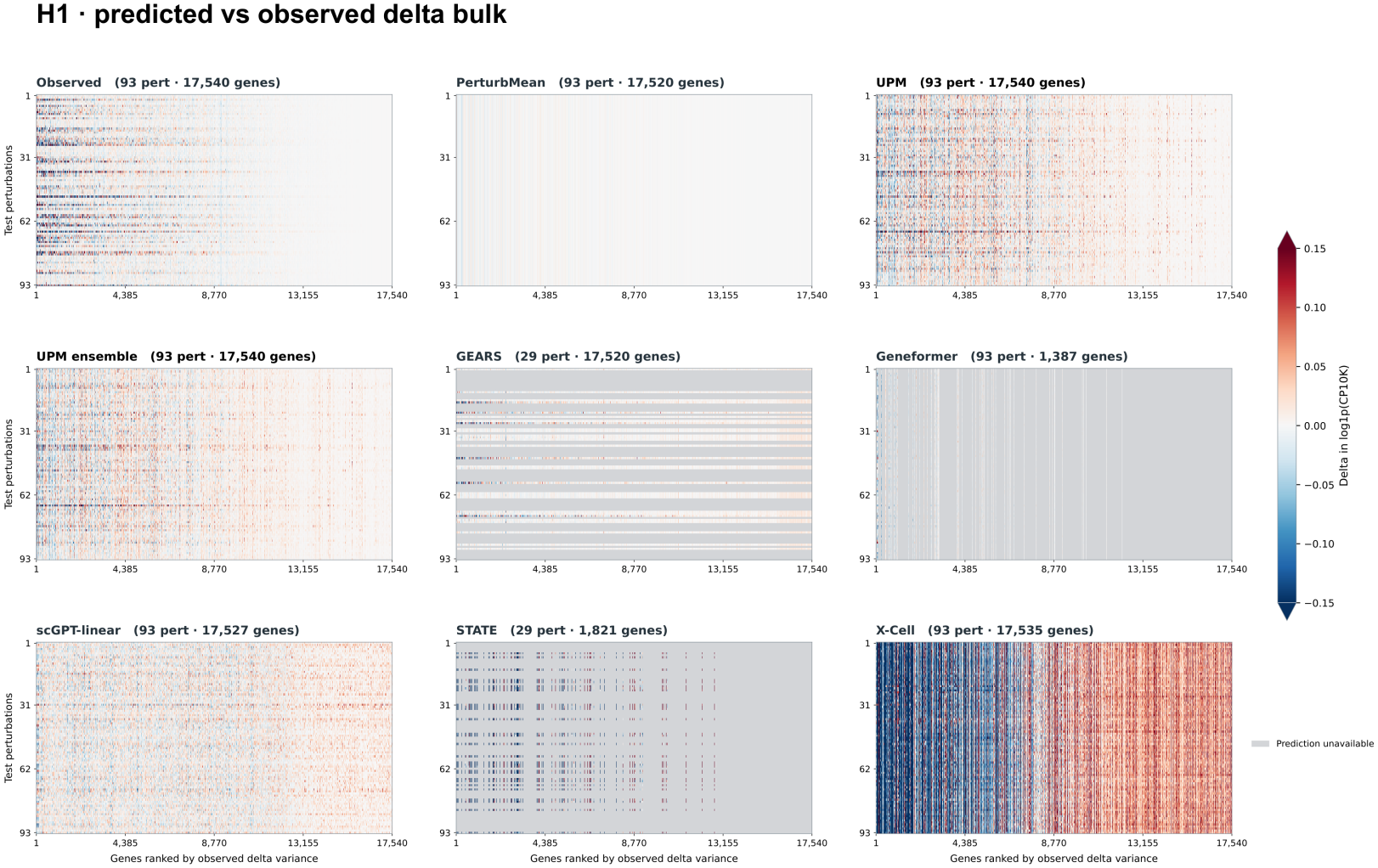
Extended mean-response heatmaps across 93 H1 perturbations. Heatmaps show control-centered mean expression across 17,540 genes, with perturbations in rows and genes in columns. Panels share row and column ordering and a symmetric color scale; gray indicates unavailable predictions. The comparison includes standalone UPM and the UPM ensemble. GEARS uses the Figure 2D checkpoint and covers 29 perturbations on 17,520 genes; Geneformer covers 1,387 genes. Processing details are provided in Appendix B.4.

**Figure S19:**
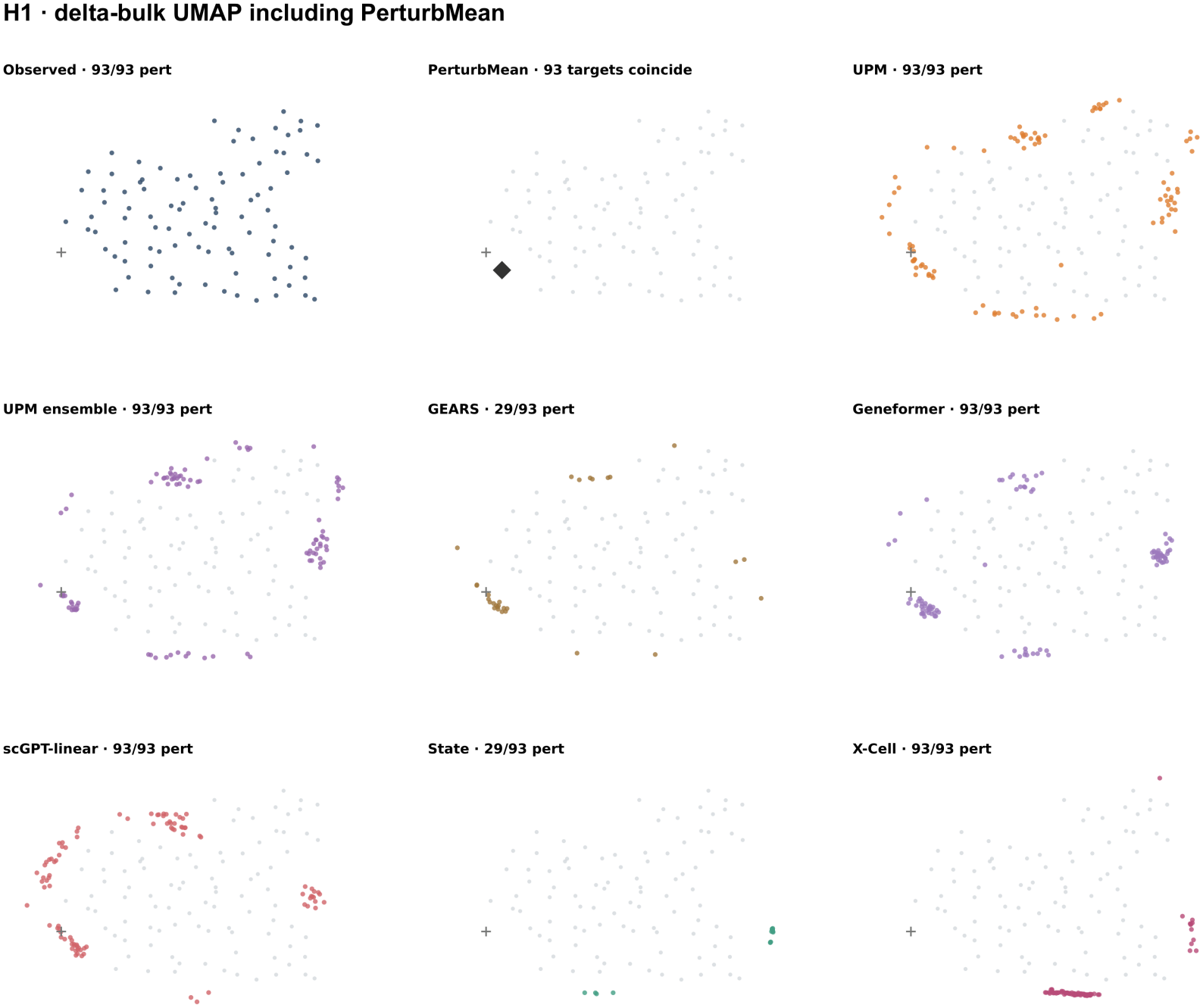
Extended mean-response UMAP across 93 H1 perturbations. Each point represents a perturbation mean response from the extended comparison in Figure S18, including standalone UPM and the UPM ensemble. PerturbMean points overlap because all perturbations receive the same predicted response.

**Figure S20:**
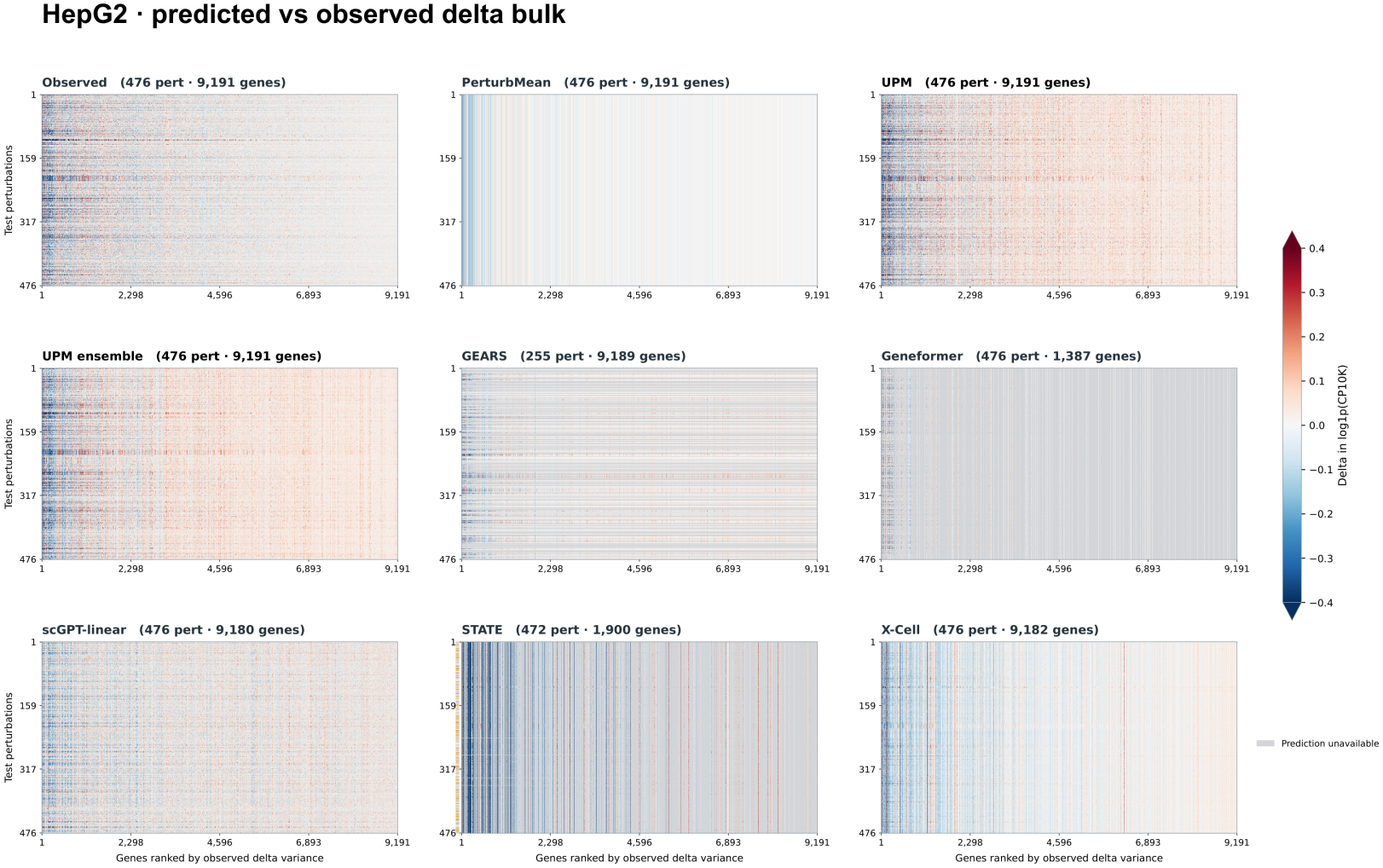
Extended mean-response heatmaps across 476 HepG2 perturbations. Heatmaps show responses across 9,191 genes using the display conventions in Figure S18, including standalone UPM and the UPM ensemble. GEARS uses the Figure 2D checkpoint and covers 255 perturbations on 9,189 genes; Geneformer covers 1,387 genes. Gray indicates unavailable predictions.

**Figure S21:**
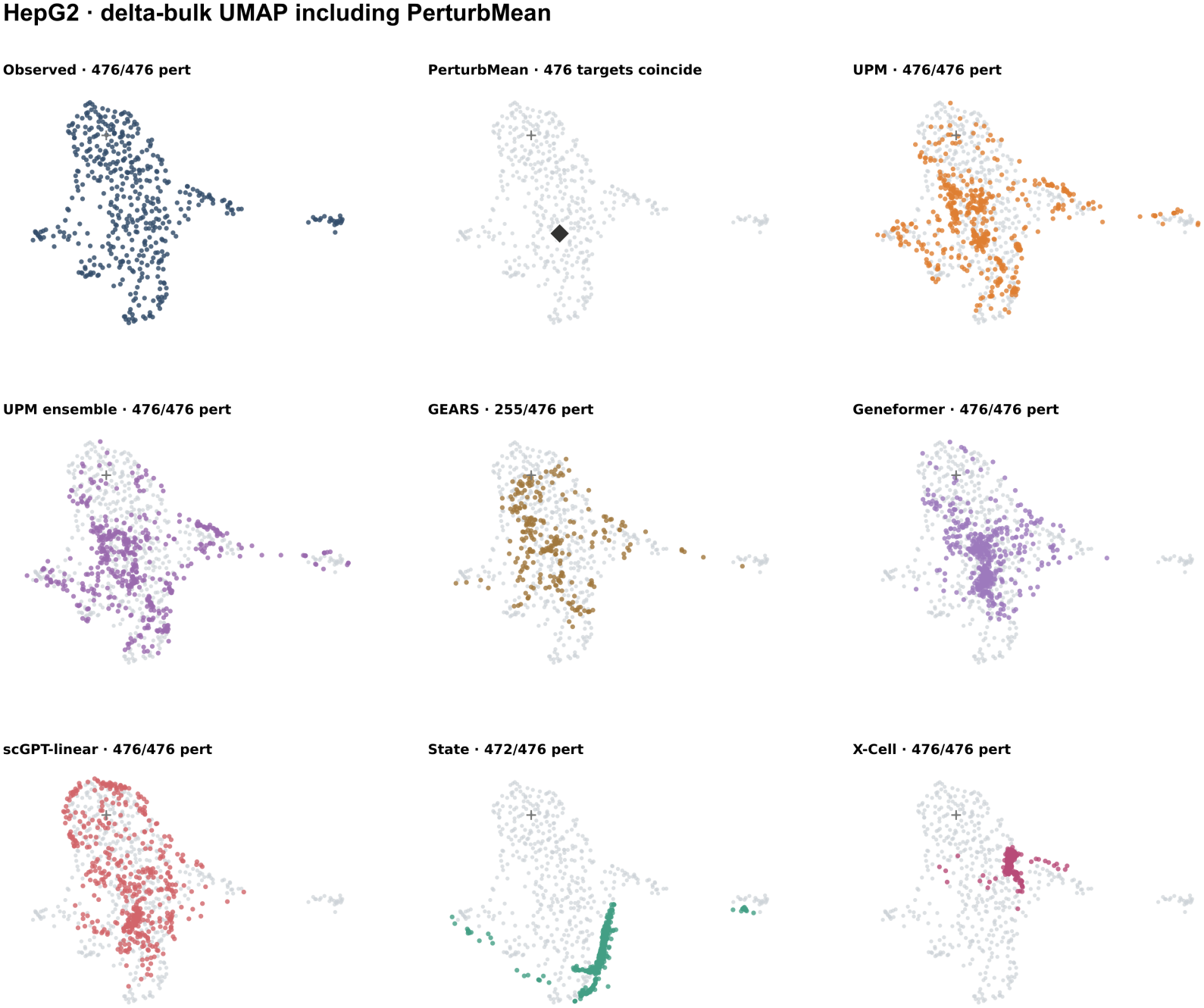
Extended mean-response UMAP across 476 HepG2 perturbations. Each point represents a perturbation mean response from the extended comparison in Figure S20, including standalone UPM and the UPM ensemble. Display conventions follow Figure S19.

**Figure S22:**
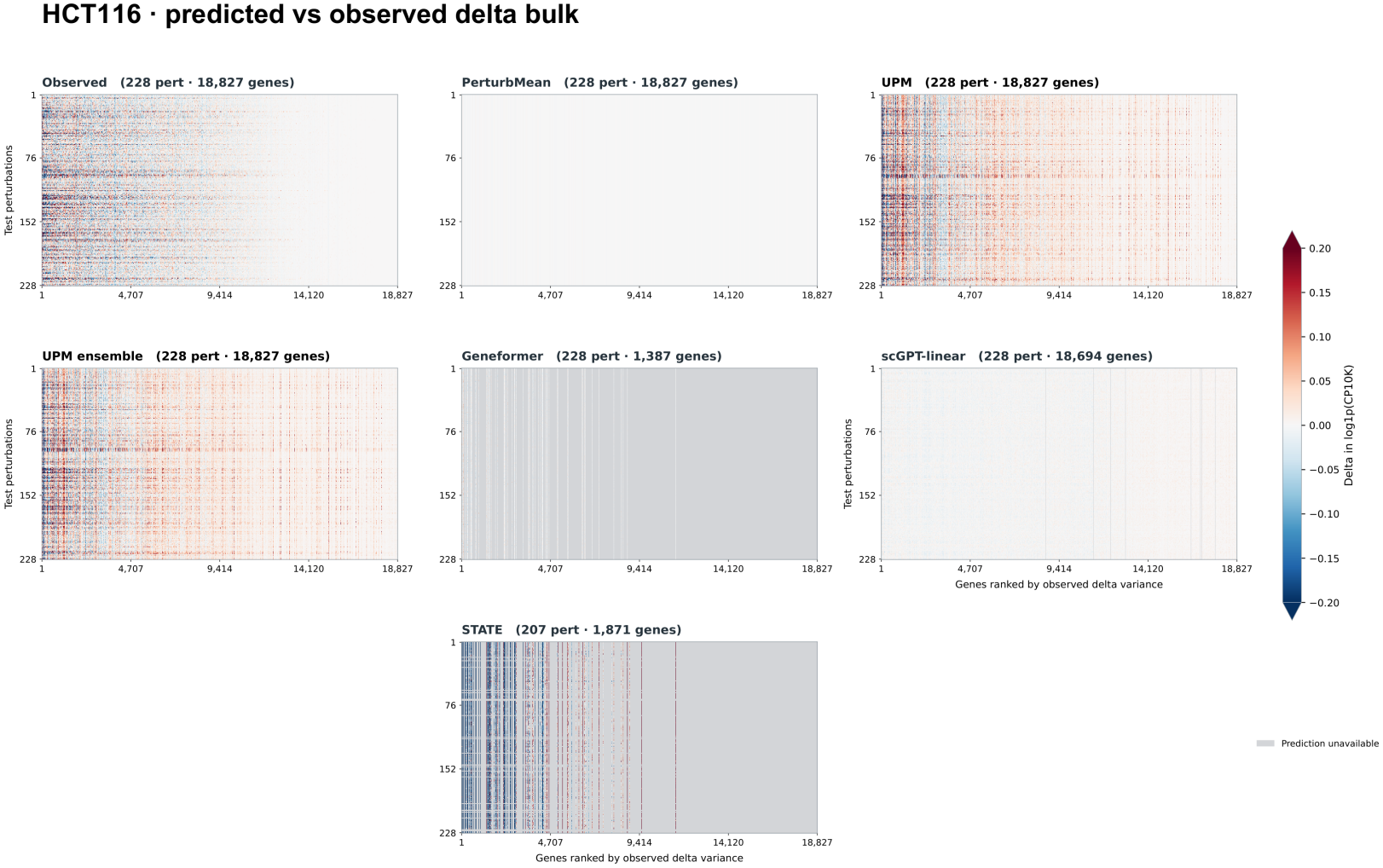
Extended mean-response heatmaps across 228 HCT116 perturbations. Heatmaps show earlier model outputs across 18,827 genes using the display conventions in Figure S18, including standalone UPM and the UPM ensemble. Geneformer covers 1,387 genes, and gray indicates unavailable predictions. GEARS and X-Cell are not included. The updated comparison in Figure 2D instead uses 207 matched perturbations and 18,907 genes.

**Figure S23:**
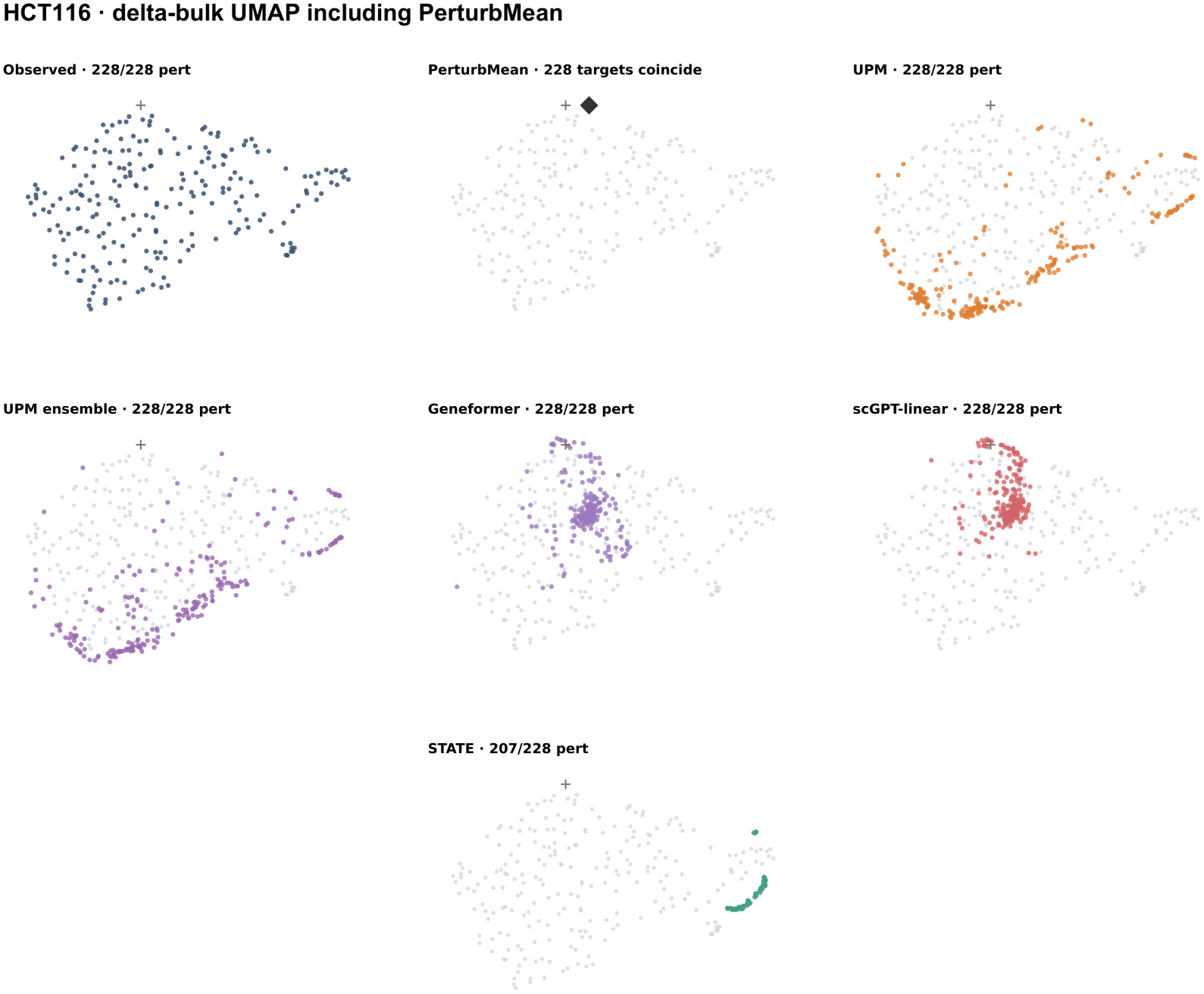
Extended mean-response UMAP across 228 HCT116 perturbations. Each point represents a perturbation mean response from the extended comparison in Figure S22, using the same 18,827-gene panel and model configurations. Display conventions follow Figure S19.

**Figure S24:**
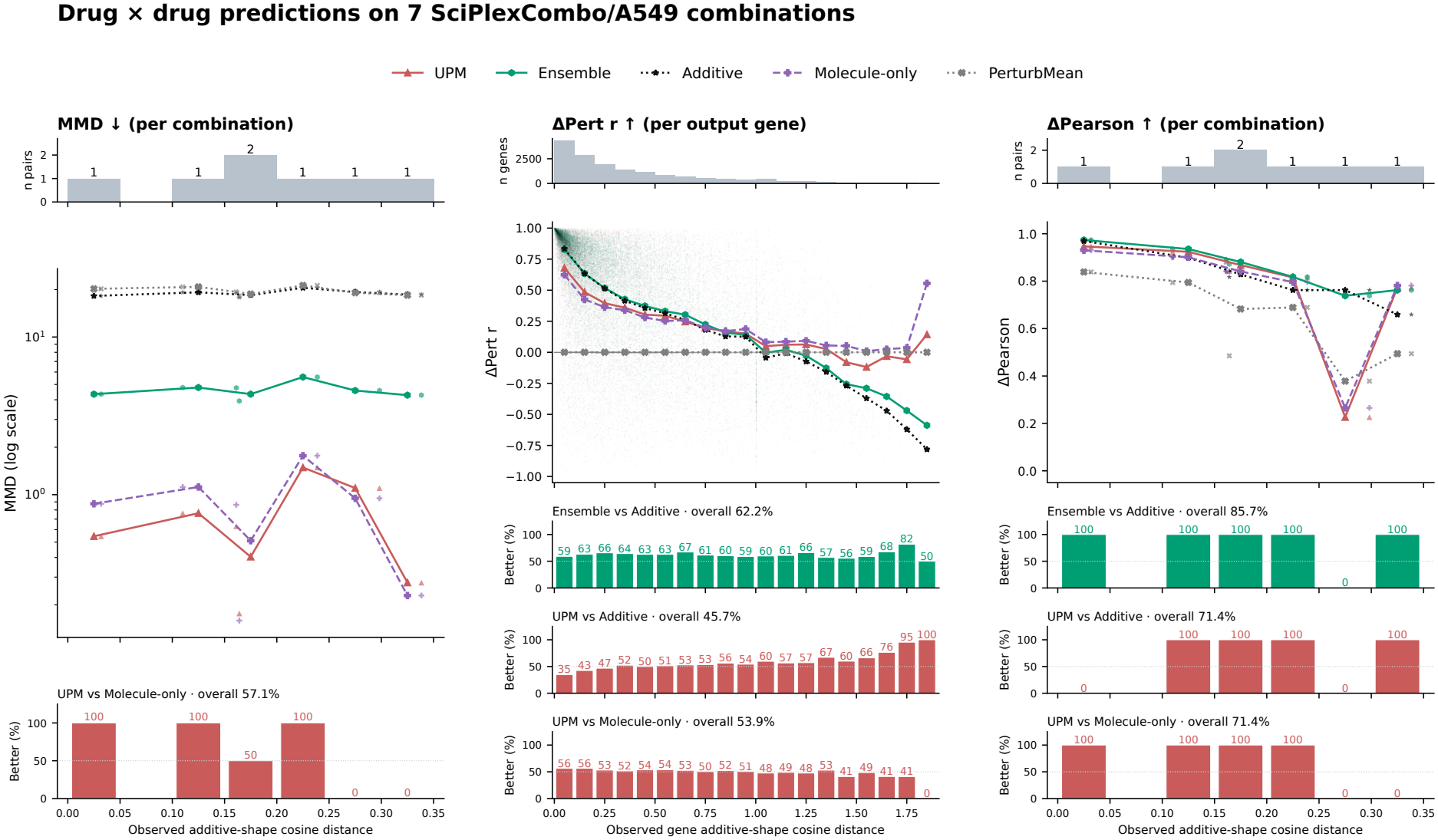
Drug-drug combination prediction across non-additivity levels. Seven held-out SciPlexCombo/A549 pairs with measured singleton responses are evaluated by MMD, gene-wise ΔPert *r* and condition-wise ΔPearson, from left to right. Non-additivity is the cosine distance between observed combination responses and Additive predictions. Histograms show combinations or genes per bin, points show individual scores, and lines connect bin means. Lower bars show pairwise win fractions, omitting comparisons uniformly at 0% or 100%. The Additive-assisted ensemble combines 50% Additive with 25% L6 and 25% L10 UPM predictions. Condition-wise trends describe these seven pairs, a subset of the 25 combinations evaluated in Figure 2B. Evaluation details are provided in Appendix B.4.1.

## References

[1] Charlotte Bunne, Yusuf Roohani, Yanay Rosen, Ankit Gupta, Xikun Zhang, Marcel Roed, Theo Alexandrov, Mohammed AlQuraishi, Patricia Brennan, Daniel B. Burkhardt, et al. How to build the virtual cell with artificial intelligence: Priorities and opportunities. Cell, 187(25):7045–7063, 2024.

[2] Yuge Ji, Mohammad Lotfollahi, F. Alexander Wolf, and Fabian J. Theis. Machine learning for perturbational single-cell omics. Cell Systems, 12(6):522–537, 2021.

[3] Jennifer E. Rood, Anna Hupalowska, and Aviv Regev.Toward a foundation model of causal cell and tissue biology with a Perturbation Cell and Tissue Atlas. Cell, 187(17):4520–4545, 2024.

[4] Atray Dixit, Oren Parnas, Biyu Li, Jenny Chen, Charles P. Fulco, Livnat Jerby-Arnon, Nemanja D. Marjanovic, Danielle Dionne, Tyler Burks, Raktima Raychowdhury, et al. Perturb-Seq: dissecting molecular circuits with scalable single-cell RNA profiling of pooled genetic screens. Cell, 167(7):1853–1866.e17, 2016.

[5] Britt Adamson, Thomas M. Norman, Marco Jost, Min Y. Cho, James K. Nuñez, Yuwen Chen, Jacqueline E. Villalta, Luke A. Gilbert, Max A. Horlbeck, Marco Y. Hein, et al. A multiplexed single-cell CRISPR screening platform enables systematic dissection of the unfolded protein response. Cell, 167(7):1867–1882.e21, 2016.

[6] Joseph M. Replogle, Reuben A. Saunders, Angela N. Pogson, Jeffrey A. Hussmann, Alexander Lenail, Alina Guna, Lauren Mascibroda, Eric J. Wagner, Karen Adelman, Gila Lithwick-Yanai, et al. Mapping information-rich genotype-phenotype landscapes with genome-scale Perturb-seq. Cell, 185(14):2559–2575.e28, 2022.

[7] Stefan Peidli, Tessa D. Green, Ciyue Shen, Torsten Gross, Joseph Min, Samuele Garda, Bo Yuan, Linus J. Schumacher, Jake P. Taylor-King, Debora S. Marks, et al. scPerturb: harmonized single-cell perturbation data. Nature Methods, 21(3):531–540, 2024.

[8] Zhiting Wei, Yiheng Wang, Yicheng Gao, Shuguang Wang, Ping Li, Duanmiao Si, Yuli Gao, Siqi Wu, Danlu Li, Kejing Dong, et al. Benchmarking algorithms for generalizable single-cell perturbation response prediction. Nature Methods, 23(2):451–464, 2026.

[9] Djordje Miladinovic, Tobias Höppe, Mathieu Chevalley, Andreas Georgiou, Lachlan Stuart, Arash Mehrjou, Marcus Bantscheff, Bernhard Schölkopf, and Patrick Schwab. In silico biological discovery with large perturbation models. Nature Computational Science, 5(11):1029–1040, 2025.

[10] Yiming Li, Min Zeng, Jun Zhu, Linjing Liu, Fang Wang, Longkai Huang, Fan Yang, Min Li, and Jianhua Yao. UniPert-G2CP bridges genetic and chemical screens from molecular representation to phenotype modeling. Cell, 189(16):4946–4963.e8, 2026.

[11] Rich Caruana. Multitask learning. Machine Learning, 28(1):41–75, 1997.

[12] Zoe Piran, Niv Cohen, Yedid Hoshen, and Mor Nitzan. Disentanglement of single-cell data with biolord. Nature Biotechnology, 42(11):1678–1683, 2024.

[13] Rishi Verma, Abhinav Adduri, Beatrice Bevilacqua, Basak Eraslan, Dave Burke, Hani Goodarzi, and Yusuf Roohani. PIE: Generalizing perturbation effects across unseen perturbations, contexts and datasets. bioRxiv, 2026.

[14] Chloe Wang, Mehran Karimzadeh, Neal G. Ravindra, Lexi R. Bounds, Nader Alerasool, Ann C. Huang, Shihao Ma, Daniel R. Gulbranson, Haotian Cui, Yongju Lee, et al. X-Cell: Scaling causal perturbation prediction across diverse cellular contexts via diffusion language models. bioRxiv, 2026.

[15] Abhinav K. Adduri, Dhruv Gautam, Beatrice Bevilacqua, Mohsen Naghipourfar, Alishba Imran, Rohan Shah, Noam Teyssier, Rishi Verma, Christopher Carpenter, Basak Eraslan, et al. Predicting cellular responses to perturbation across diverse contexts with State. Cell, 189(19):5914–5931.e20, 2026.

[16] Mingze Dong, Abhinav Adduri, Dhruv Gautam, Lujing Wu, Courtney Kernick, Mary Margaret Coons, Yi-Chen Chih, Christopher Carpenter, Rohan Shah, Chiara Ricci-Tam, et al. Stack: In-context learning of single-cell biology. bioRxiv, 2026.

[17] Hyun Young Kim, Sara Brin Rosenthal, Xiao Liu, Charlene Miciano, Xiaomeng Hou, Michael Miller, Justin Buchanan, Olivier B. Poirion, Daisy Chilin-Fuentes, Cuijuan Han, et al. Multi-modal analysis of human hepatic stellate cells identifies novel therapeutic targets for metabolic dysfunction-associated steatotic liver disease. Journal of Hepatology, 82(5):882–897, 2025.

[18] Xiao Liu, Jun Xu, Sara Rosenthal, Ling-juan Zhang, Ryan McCubbin, Nairika Meshgin, Linshan Shang, Yukinori Koyama, Hsiao-Yen Ma, Sonia Sharma, et al. Identification of lineage-specific transcription factors that prevent activation of hepatic stellate cells and promote fibrosis resolution. Gastroenterology, 158(6):1728–1744.e14, 2020.

[19] Ajay Nadig, Joseph M. Replogle, Angela N. Pogson, Mukundh Murthy, Steven A. McCarroll, Jonathan S. Weissman, Elise B. Robinson, and Luke J. O’Connor. Transcriptome-wide analysis of differential expression in perturbation atlases. Nature Genetics, 57(5):1228–1237, 2025.

[20] Thomas M. Norman, Max A. Horlbeck, Joseph M. Replogle, Alex Y. Ge, Albert Xu, Marco Jost, Luke A. Gilbert, and Jonathan S. Weissman. Exploring genetic interaction manifolds constructed from rich single-cell phenotypes. Science, 365(6455):786–793, 2019.

[21] Yusuf H. Roohani, Tony J. Hua, Po-Yuan Tung, Lexi R. Bounds, Feiqiao B. Yu, Alexander Dobin, Noam Teyssier, Abhinav Adduri, Alden Woodrow, Brian S. Plosky, et al. Virtual Cell Challenge: Toward a Turing test for the virtual cell. Cell, 188(13):3370–3374, 2025.

[22] Ann C. Huang, Tsung-Han S. Hsieh, Jiang Zhu, Jackson Michuda, Ashton Teng, Soohong Kim, Elizabeth M. Rumsey, Sharon K. Lam, Ikenna Anigbogu, Philip Wright, et al. X-Atlas/Orion: Genome-wide Perturb-seq datasets via a scalable Fix-Cryopreserve platform for training dose-dependent biological foundation models. bioRxiv, 2025.

[23] Sanjay R. Srivatsan, José L. McFaline-Figueroa, Vijay Ramani, Lauren Saunders, Junyue Cao, Jonathan Packer, Hannah A. Pliner, Dana L. Jackson, Riza M. Daza, Lena Christiansen, et al. Massively multiplex chemical transcriptomics at single-cell resolution. Science, 367(6473):45–51, 2020.

[24] Mohammad Lotfollahi, Anna Klimovskaia Susmelj, Carlo De Donno, Leon Hetzel, Yuge Ji, Ignacio L. Ibarra, Sanjay R. Srivatsan, Mohsen Naghipourfar, Riza M. Daza, Beth Martin, et al. Predicting cellular responses to complex perturbations in high-throughput screens. Molecular Systems Biology, 19(6):e11517, 2023.

[25] Jesse Zhang, Airol A. Ubas, Valentine Svensson, Nicole Thomas, Vishvak Subramanyam, Christopher Carpenter, Aidan Winters, Richard de Borja, Noam Teyssier, Neha Thakar, et al. Tahoe-100M: Mapping drug-induced molecular phenotypes at single-cell resolution. Cell, 189(19):5945–5961.e8, 2026.

[26] Lukas Oesinghaus, Sören Becker, Larsen Vornholz, Efthymia Papalexi, Joey Pangallo, Amir Ali Moinfar, Jenni Liu, Alyssa La Fleur, Maiia Shulman, Simone Marrujo, et al. A single-cell cytokine dictionary of human peripheral blood. bioRxiv, 2025.

[27] Yusuf Roohani, Kexin Huang, and Jure Leskovec. Predicting transcriptional outcomes of novel multigene perturbations with GEARS. Nature Biotechnology, 42(6):927–935, 2024.

[28] Christina V. Theodoris, Ling Xiao, Anant Chopra, Mark D. Chaffin, Zeina R. Al Sayed, Matthew C. Hill, Helene Mantineo, Elizabeth M. Brydon, Zexian Zeng, X. Shirley Liu, et al. Transfer learning enables predictions in network biology. Nature, 618(7965):616–624, 2023.

[29] Haotian Cui, Chloe Wang, Hassaan Maan, Kuan Pang, Fengning Luo, Nan Duan, and Bo Wang. scGPT: toward building a foundation model for single-cell multi-omics using generative AI. Nature Methods, 21(8):1470–1480, 2024.

[30] José L. McFaline-Figueroa, Sanjay Srivatsan, Andrew J. Hill, Molly Gasperini, Dana L. Jackson, Lauren Saunders, Silvia Domcke, Samuel G. Regalado, Paul Lazarchuck, Sarai Alvarez, et al. Multiplex single-cell chemical genomics reveals the kinase dependence of the response to targeted therapy. Cell Genomics, 4(2):100487, 2024.

[31] Henry E. Miller, Gabriel M. Mejia, Francis J. A. Leblanc, Brendan Swain, Bo Wang, and Lucas Paulo de Lima Camillo. Deep learning perturbation models can outperform baselines on calibrated metrics. Nature Biotechnology, 2026.

[32] Arc Institute. Virtual Cell Challenge: Evaluation, 2026. https://virtualcellchallenge.org/evaluation.

[33] Mohammad Lotfollahi, F. Alexander Wolf, and Fabian J. Theis. scGen predicts single-cell perturbation responses. Nature Methods, 16(8):715–721, 2019.

[34] Leon Hetzel, Simon Boehm, Niki Kilbertus, Stephan Günnemann, Mohammad Lotfollahi, and Fabian Theis. Predicting cellular responses to novel drug perturbations at a single-cell resolution. In Advances in Neural Information Processing Systems, volume 35, pages 26711–26722. Curran Associates, Inc., 2022.

[35] Chen Li, Lei Wei, and Xuegong Zhang. Unified multimodal learning enables generalized cellular response prediction to diverse perturbations. bioRxiv, 2026.

[36] Xiaoning Qi, Lianhe Zhao, Chenyu Tian, Yueyue Li, Zhen-Lin Chen, Peipei Huo, Runsheng Chen, Xiaodong Liu, Baoping Wan, Shengyong Yang, et al. Predicting transcriptional responses to novel chemical perturbations using deep generative model for drug discovery. Nature Communications, 15(1):9256, 2024.

[37] Dominik Klein, Jonas Simon Fleck, Daniil Bobrovskiy, Lea Zimmermann, Sören Becker, Alessandro Palma, Leander Dony, Alejandro Tejada-Lapuerta, Guillaume Huguet, Hsiu-Chuan Lin, et al. CellFlow enables generative single-cell phenotype modeling with flow matching. bioRxiv, 2025.

[38] Zeming Lin, Halil Akin, Roshan Rao, Brian Hie, Zhongkai Zhu, Wenting Lu, Nikita Smetanin, Robert Verkuil, Ori Kabeli, Yaniv Shmueli, et al. Evolutionary-scale prediction of atomic-level protein structure with a language model. Science, 379(6637):1123–1130, 2023.

[39] Charlotte Bunne, Stefan G. Stark, Gabriele Gut, Jacobo Sarabia del Castillo, Mitch Levesque, Kjong-Van Lehmann, Lucas Pelkmans, Andreas Krause, and Gunnar Rätsch. Learning single-cell perturbation responses using neural optimal transport. Nature Methods, 20(11):1759–1768, 2023.

[40] Chenglei Yu, Chuanrui Wang, Bangyan Liao, and Tailin Wu. scDFM: Distributional flow matching model for robust single-cell perturbation prediction. arXiv preprint arXiv:2602.07103, 2026.

[41] Siyu He, Yuefei Zhu, Daniel Naveed Tavakol, Haotian Ye, Yeh-Hsing Lao, Zixian Zhu, Cong Xu, Shradha Chauhan, Guy Garty, Raju Tomer, et al. Squidiff: predicting cellular development and responses to perturbations using a diffusion model. Nature Methods, 23(1):65–77, 2026.

[42] Han Zhang, Guo-Hua Yuan, Chaohao Yuan, Tingyang Xu, Tian Bian, Hong Cheng, Wenbing Huang, Deli Zhao, and Yu Rong. Lingshu-Cell: A generative cellular world model for transcriptome modeling toward virtual cells. arXiv preprint arXiv:2603.25240, 2026.

[43] Johann Wenckstern, Eeshaan Jain, Benedikt von Querfurth, Yexiang Cheng, Kiril Vasilev, Matteo Pariset, Phil F. Cheng, Petros Liakopoulos, Olivier Michielin, Andreas Wicki, et al. The Virtual Tissues foundation model resolves spatial proteomics across scales. Nature, 657(8132):744–754, 2026.

[44] Yunrui Lu, Willis Zhang, Yu-Jen Chen, Jiayi Yin, Lydia Chen, Kevin Fleisher, James Gornet, Rex Liu, Zitong Jerry Wang, Yan Poon, et al. A generative virtual tissue model enables computational design of therapeutic perturbation strategies. bioRxiv, 2026.

[45] Constantin Ahlmann-Eltze, Wolfgang Huber, and Simon Anders. Deep-learning-based gene perturbation effect prediction does not yet outperform simple linear baselines. Nature Methods, 22(8):1657–1661, 2025.

[46] Daniel R. Wong, Abby S. Hill, and Rob Moccia. Simple controls exceed best deep learning algorithms and reveal foundation model effectiveness for predicting genetic perturbations. Bioinformatics, 41(6):btaf317, 2025.

[47] Ramon Viñas Torné, Maciej Wiatrak, Zoe Piran, Shuyang Fan, Liangze Jiang, Sarah A. Teichmann, Mor Nitzan, and Maria Brbić. Systema: a framework for evaluating genetic perturbation response prediction beyond systematic variation. Nature Biotechnology, 44(6):1050–1059, 2026.

[48] Abhinav Adduri, Dave Burke, Joseph Caputo, Paul Datlinger, Alexander Dobin, Basak Eraslan, Luke Gilbert, Patrick Hsu, Ben Hsu, Tony Hua, et al. Virtual Cell Challenge 2026: Benchmarking zero-shot generalization across cellular contexts. Cell, 189(19):5827–5830, 2026.

[49] David Rogers and Mathew Hahn. Extended-connectivity fingerprints. Journal of Chemical Information and Modeling, 50(5):742–754, 2010.

[50] Arthur Gretton, Karsten M. Borgwardt, Malte J. Rasch, Bernhard Schölkopf, and Alexander Smola. A kernel two-sample test. Journal of Machine Learning Research, 13(25):723–773, 2012.

